# COPII-coated membranes function as transport carriers of intracellular procollagen-1

**DOI:** 10.1101/110700

**Authors:** Amita Gorur, Lin Yuan, Samuel J Kenny, Satoshi Baba, Ke Xu, Randy Schekman

**Affiliations:** Department of Molecular and Cell Biology and Howard Hughes Medical Institute, University of California, Berkeley, CA 94720.; Department of Chemistry, University of California, Berkeley, CA 94720.

## Abstract

The coat protein complex II (COPII) is essential for the secretion of large cargo, such as the 300 nm precursor fibrils of procollagen I (PC1). Previous work has shown that the CUL3-KLHL12 complex increases the size of COPII vesicles to over 300 nm in diameter and accelerates the secretion of PC1; however, the role of large COPII vesicles as PC1 transport carriers was not unambiguously demonstrated. In this study, using stochastic optical reconstruction microscopy (STORM), correlated light electron microscopy (CLEM), and live cell imaging we report the existence of mobile COPII-coated vesicles that completely encapsulate the cargo PC1 and are physically separated from ER. We have also developed a cell-free COPII vesicle budding reaction that reconstitutes the capture of PC1 into large COPII vesicles. This process requires COPII proteins and the GTPase activity of the COPII subunit SAR1. We conclude from *in vivo* and *in vitro* evidence that large COPII vesicles are *bona fide* carriers of PC1.

**Summary:** COPII may play a direct or indirect role in the traffic of large protein complexes such as procollagen. Using high resolution imaging techniques in intact cells and in vitro reconstituted vesicles, Gorur et al. show that COPII coated vesicles carry procollagen1.

## Introduction

As an essential step in conventional protein secretion, coat protein complex II (COPII) mediates vesicular transport from the ER to the Golgi apparatus in eukaryotes. The GTPase SAR1, inner coat proteins SEC23/SEC24, and outer coat proteins SEC13/SEC31 are five cytosolic components of the COPII complex, and they are sufficient to generate COPII-coated vesicles from synthetic liposomes (Matsuoka et al., 1998; Kim et al., 2005). COPII vesicles were observed by EM to be about 60-80 nm in diameter which potentially limits the transport of large cargos such as the 300 nm long procollagen I (PC1) rigid rod (Barlowe et al., 1994; Bächinger et al., 1982; Kim et al., 2005; Noble et al., 2013). However, human genetic evidence showed that COPII is required to secrete procollagens (PC). Mutations in genes that code for the human COPII paralogs SEC23A and SEC24D were identified to cause the genetic diseases cranio-lenticulo-sutural dysplasia and osteogenesis imperfecta and their characteristic collagen deposition defects during development (Boyadjiev et al., 2006; Kim et al., 2012; Garbes et al., 2015).

The aforementioned requirement for COPII to secrete PC has been independently demonstrated in multiple model systems. Mutation of the *sec-23* gene in *Caenorhabditis elegans* disrupts collagen secretion and leads to aberrant cuticle, dissociated hypodermal cells and late embryonic lethality (Roberts et al., 2003). In *Drosophila melanogaster,* tissue-specific knockdown of Sar1 or Sec23 in the collagen-secreting fat body cells leads to intracellular accumulation of collagen and cell lethality (Pastor-Pareja and Xu, 2011). The zebrafish mutants *crusher* and *bulldog* resulted from mutations in *sec23a* and *sec24d* genes respectively, and their chondrocytes retain procollagen II in the ER. These mutants also show defects during craniofacial development with phenotypes reminiscent of the human disease cranio-lenticulo-sutural dysplasia (Lang et al., 2006; Sarmah et al., 2010). Sec23A null mice are embryonically lethal and skin fibroblasts accumulate ER-localized collagen I and III (Zhu et al., 2015). Knock down of SEC13 in primary human dermal fibroblasts also selectively blocks PC1 secretion (Townley et al., 2008). Hence, the requirement for COPII in the ER exit of PC is evolutionarily conserved in metazoans.

The necessary role of COPII in large cargo secretion is further supported by the discovery of a large transmembrane protein TANGO1 (MIA3), which has been shown to have a general role in the secretion of large cargos, including many members of the collagen family, laminin, and large lipoprotein complexes such as pre-chylomicrons (Saito et al., 2009; Wilson et al., 2011; Petley-Ragan et al., 2016; Santos et al., 2016). The luminal Src homology 3 domain of TANGO1 interacts with the PC-specific chaperone HSP47 to recognize a broad range of PC isoforms (Saito et al., 2009; Ishikawa et al., 2016). The cytosolic side of TANGO1 was shown to interact with multiple COPII components: its proline-rich domain binds to the inner COPII coat protein SEC23 directly, and its second coiled-coil domain recruits cTAGE5, a spliced variant of a TANGO1 isoform, which binds SEC12, an initiating factor of COPII assembly (Saito et al., 2009, 2011, 2014;Ma and Goldberg, 2016). Therefore, TANGO1 plays an important role in coordinating large cargo sensing and COPII recruitment, which further supports the involvement of COPII in large cargo secretion.

Although the requirement for COPII to export the large cargo PC out of the ER is clear, the precise role that COPII plays in this process is poorly understood. A conventional model was proposed in which COPII concentrates large cargos at ER exit sites (ERES), orchestrates the packaging of large cargos into vesicles and the formation of vesicles with structured coats (Fromme and Schekman, 2005). An alternative model suggests that COPII only functions to concentrate large cargos and other factors required for the ER export at ERES, but large cargos exit the ER in carriers not coated with COPII proteins (Mironov et al., 2003; Siddiqi et al., 2003; Siddiqi et al., 2010).

The conventional model is paradoxical unless a cellular mechanism exists to increase the size of COPII-coated vesicles. The Rape and Schekman laboratories previously reported that the E3 ubiquitin ligase, CUL3, and one of its substrate adaptors, KLHL12, regulate the size of COPII-coated vesicles and collagen I and IV secretion (Jin et al., 2012). Overexpression of KLHL12 induces the formation of large COPII structures that are decorated with KLHL12. Most of these structures are more than 300 nm in diameter and large enough to accommodate cargo of the size of PC1. Unfortunately, the coincident localization of PC1 in these large carriers was not clearly elucidated in this work. Here, we re-examined these large COPII structures and showed that they are *bona fide* membranous carriers of PC1 in cells. Moreover, we reconstituted PC1 capture into vesicles formed in a cell-free reaction. These carriers were isolated and visualized as large COPII-coated vesicles.

## Results

### Large COPII vesicles co-localize with PC1

The ubiquitin ligase CUL3 and its substrate adaptor KLHL12 were shown to regulate collagen secretion and the size of COPII vesicles, possibly through the monoubiquitylation of the COPII protein SEC31A (Jin et al., 2012). To study whether the large COPII vesicles observed after overexpression of KLHL12 are collagen carriers, we engineered human fibrosarcoma cells (KI6) to stably overexpress the human pro-α1(I) collagen and inducible KLHL12 under doxycycline-controlled transcriptional activation. KI6 cells synthesized PC1 in contrast to the parent cell line as confirmed by an immunofluorescence signal for PC1 (Fig. S1A), and immuno-gold labeled PC1 in the ER and large vesicles (Fig. S1B). We also observed accelerated ER to Golgi trafficking of PC1 in KI6 cells after KLHl12 overexpression was induced for 7.5 h, which was consistent with our previous report (Fig. S1C, Jin et al., 2012). KI6 cells secreted PC1 through the conventional secretory pathway, shown by the inhibitory effect of brefeldin A (BFA) on the export of PC1 into the culture medium (Fig. S1D). Because TANGO1 KO mice are defective in collagen I secretion, we knocked down TANGO1 in KI6 cells and observed a decreased level of PC1 secretion as detected by immunoblots of the culture medium with a corresponding intracellular accumulation of PC1 detected in cell lysates (Fig. S1E, Wilson et al., 2011).

In our previous study (Jin et al., 2012) using a polyclonal antibody (LF-67) raised against synthetic human α1(I) collagen C-telopeptide (Bernstein et al., 1996), we observed partial colocalization between PC1 and KLHL12 but could not unambiguously document that the large COPII vesicles, generated by overexpression of KLHL12, carried PC1. In contrast, with the use of two monoclonal antibodies raised against N or C propeptides of PC1 (Fig. S2A, QED Biosciences; Personal communication, Foellmer et al., 1983), we observed clear co-localization of PC1 and KLHL12 in KI6 cells after 7.5 h of induction (Fig. 1A, Fig. S2B). Puncta positive for both PC1 and KLHL12 also co-localized with SEC31A, a subunit of the outer coat of COPII vesicles (Fig. 1B).

**Figure 1.**
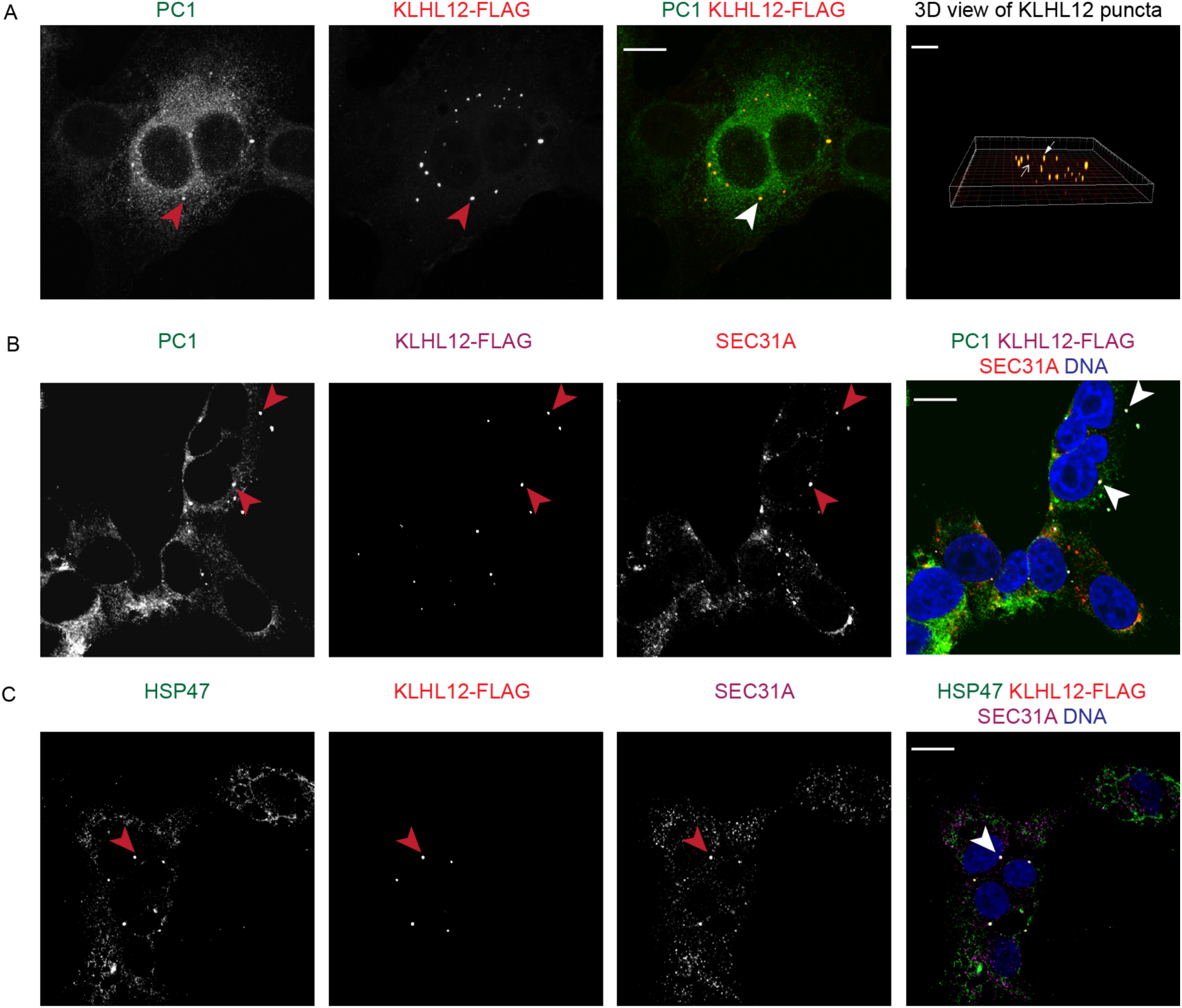
Large COPII vesicles co-localize with PC1. (A) PC1 (green) and KLHL12 (red) co-localize in KI6 cells that were treated with doxycycline for 7.5 h to induce the expression of KLHL12. Images shown are average intensity projected views of a Z stack shown for each and merged channel. 3D view of KLHL12 puncta was obtained from the Z stack using the Spot feature in Imaris 8.1.2. An example of co-localizing “yellow” puncta are indicated by a closed arrow, and KLHL12 positive puncta not co-localizing with PC1 are indicated by an open arrow. Total number of KLHL12 positive puncta determined in the Z stack was 27 and 78% of these spots were positive for PC1 calculated using CoLoc feature in Imaris 8.1.2. (B) PC1 (green), KLHL12 (magenta) and SEC31A (red) co-localize in KI6 cells (closed arrows). DAPI is shown in blue. Half of the total number of KLHL12 positive puncta (n=14) were positive for PC1 and SEC31A. (C) HSP47 (green), KLHL12 (red) and SEC31A (magenta) co-localize in KI6 cells (close arrow). Bars: 10 μm. n=3.

We reasoned that if large COPII vesicles are functional secretory transporters of collagen, they should also contain HSP47, a collagen-specific chaperone. HSP47 binds to the triple helical region of PC in the ER and promotes the correct folding of trimerized PC (Nagai et al., 2000; Tasab et al., 2000; Ono et al., 2012; Wildmer et al., 2012). HSP47 accompanies PC to the ER Golgi Intermediate Compartment (ERGIC) or cis-Golgi membrane where it dissociates because of a lower pH, after which it is recycled back to the ER via its C-terminal RDEL sequence (Satoh et al., 1996; Oecal et al., 2016). We found that HSP47 co-localized with the large COPII vesicles as visualized by triple immunofluorescence labeling of HSP47, SEC31A and KLHL12 in KI6 cells, consistent with a role for the large COPII vesicles as secretory carriers of PC1 (Fig. 1C).

### Large COPII vesicles are hollow membranous containers

Given that conventional confocal microscopy images revealed co-localized diffraction-limited puncta with no discernible morphological details, we sought to resolve these structures using stochastic optical reconstruction microscopy (STORM) (Rust *et al.* 2006; Huang *et al.* 2008), a type of super-resolution microscopy, and correlative light and electron microscopy (CLEM). With 3D STORM analysis, we observed large, greater than 300nm, hollow, cage-like COPII structures using a SEC31A antibody and a secondary antibody conjugated with Alexa Fluor 647 fluorophore in KI6 cells induced for the overexpression of KLHL12 (Fig. 2A). A virtual Z-stack of a single structure confirmed the cage-like COPII protein localization completely surrounding a cavity in three dimensions. Similar analysis was conducted on vesicles immunolabeled with FLAG antibody, which targeted the overexpressed KLHL12-FLAG. Virtual cross-sections of the structure in XY, XZ and YZ dimensions revealed a hollow compartment surrounded by a protein coat, presumably made up of the inner and outer COPII coat protein shells (Fig. 2B).

**Figure 2.**
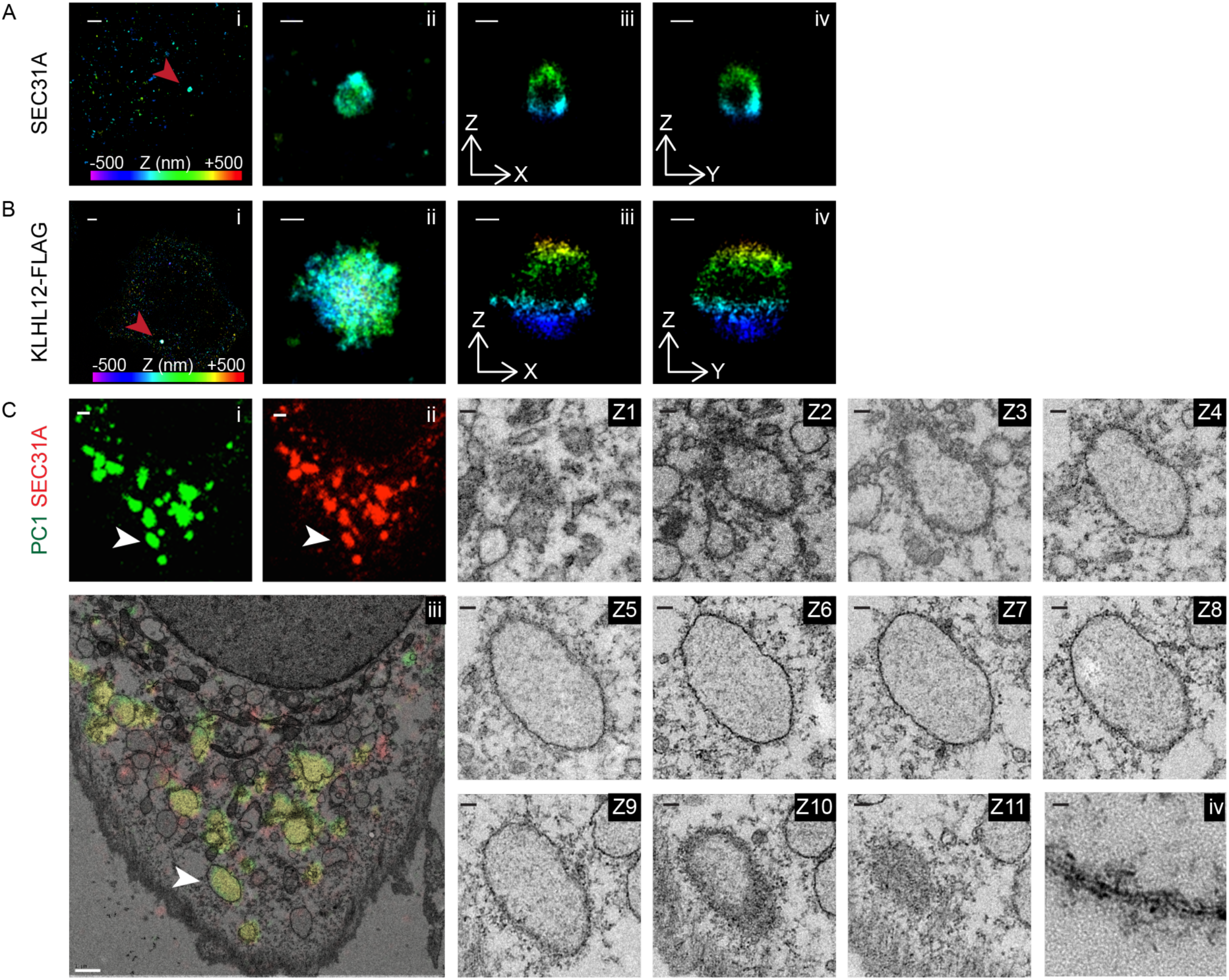
Large COPII vesicles are hollow membranous containers. 3D STORM images of large SEC31A (A) or KLHL12-FLAG (B) puncta in KI6 cells after 7.5 h induction. Position in depth (Z axis) is represented by color according to the color scale bar in (Ai and Bi). (i) Overview of an area of the cell imaged by STORM: SEC31A (A) or FLAG (B). (ii) Magnified maximum XY-projection of a large SEC31A (A) or FLAG (B) structure. Virtual cross-sections of this structure in XZ (iii) and YZ (iv) planes reveal a hollow compartment encapsulated by SEC31A (A) or KLHL12-FLAG (B). (C) CLEM resolved overlapping immunofluorescent (i) PC1 (green) and (ii) SEC31A (red) puncta as single membrane bound compartments in cultured Saos-2 cells expressing endogenous levels of PC1 and KLHL12. (iii) A representative cell showing co-localizing puncta (indicated by white arrow) was processed for CLEM. Image shows overlay of the merged fluorescent channels (green and red) on a thin section (70nm) TEM image (Z1-11) Serial sections of 70 nm thicknesses through structure of interest. (iv) Magnified view of lipid bilayer from an area of the single membrane enclosing the organelle. n=2. Scale bars in (A i): 1 μm; (B i): 1 μm; (A ii-iv): 200 nm, (B ii-iv): 200 nm; (C i-iii): 1 μm; Z1 —Z11: 100 nm, (Civ): 10 nm

We observed a similar morphological feature by CLEM on regions of co-localized endogenous PC1 and SEC31A without KLHL12 overexpression in human osteosarcoma Saos-2 cells, a cell line that secretes endogenous PC1 (Fig. 2C, Pautke et al., 2004). Correlative serial thin-section EM revealed that fluorescent puncta containing co-localized PC1 and SEC31A were large, single membrane-bounded compartments and were not clusters of small vesicles. These correlated structures were spherical to ovoid in nature and ranged from 350 μm to 1.7 μm in diameter, decorated with what may be remnants of coat protein. Taken together, the diffraction-limited PC1 and COPII co-localized puncta were resolved by STORM and CLEM to be large protein-coated single membrane-bounded containers.

### PC1 is completely encapsulated in large COPII-coated membranes

We next sought to dissect the precise localization of PC1 with respect to the COPII coat within puncta that showed co-localization of the two markers. To achieve this, we performed dual and triple color 3D STORM imaging on SEC31A/PC1 co-localizing puncta from cells with overexpressed and endogenous levels of KLHL12. Large COPII cages with hollow cavities were observed in KI6 cells (Fig. 3Aix), consistent with our observation using single color STORM (Fig. 2Aiii,iv), and PC1 was resolved to be inside of the hollow cavities, entirely encapsulated by the COPII cage (Fig. 3Av-x). This was also evident in KLHL12/SEC31A/PC1 co-localizing puncta in Saos-2 cells (Fig. 3B). The appearance of confined labeling of the PC1 fiber in relation to the expansive COPII coat may reflect a restricted orientation of the fiber within the vesicle, or steric hindrance of the PC1 monoclonal antibody by the polyclonal antibody used to label the coat. In contrast, we did not observe PC1 in canonical COPII cages ranging in size from 80-100 nm (Fig. 3A ii-iv). Interestingly, we also observed puncta that appeared to have the COPII coat only partially enveloping PC1 possibly representing an intermediate in the shedding of COPII subunits, or a nascent budding event at the ERES (Fig. 3B, Fig. S3).

**Figure 3.**
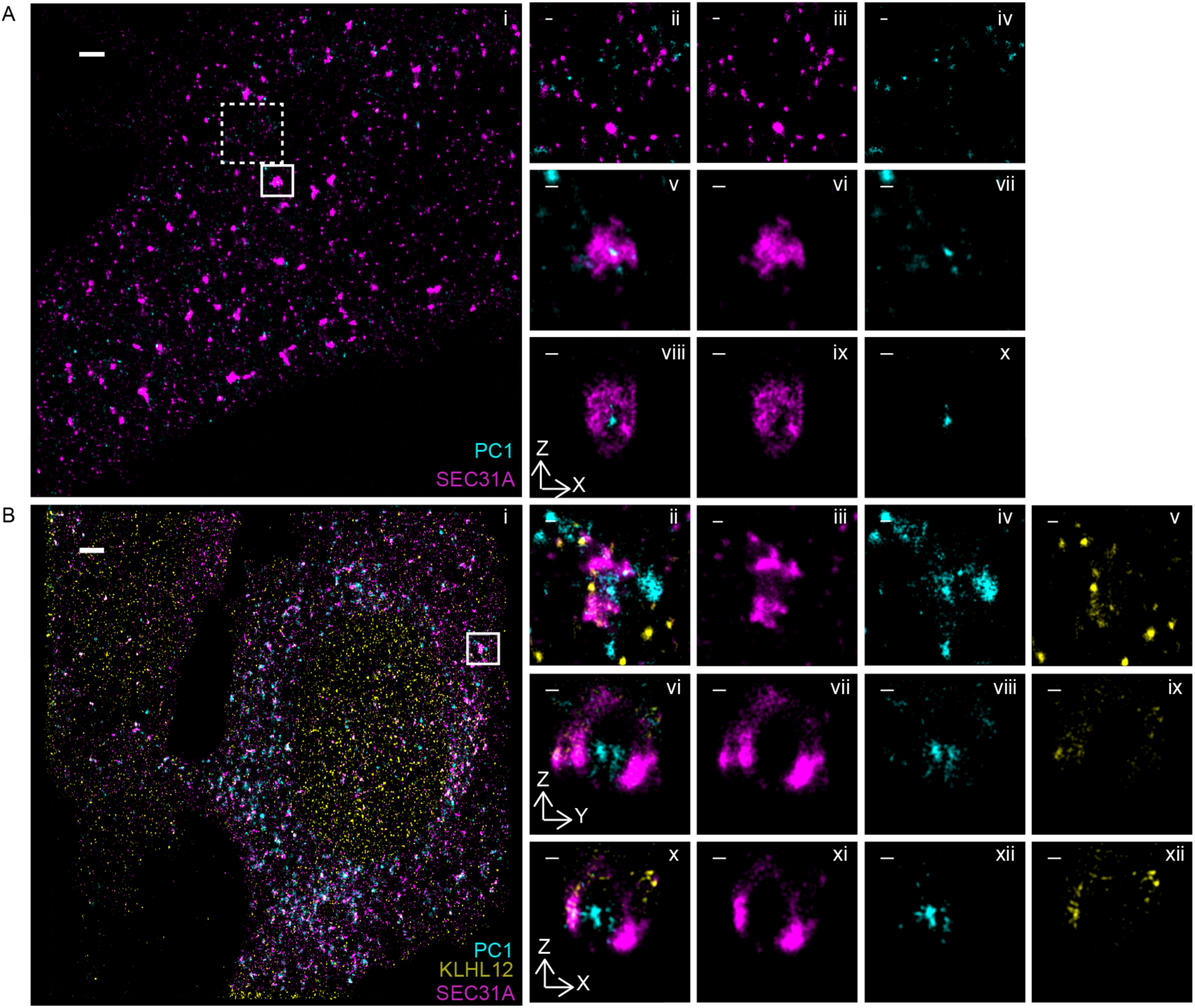
PC1 is completely encapsulated in large COPII-coated vesicles. (A) Two-color 3D STORM revealed PC1 (cyan) residing inside of large SEC31A (magenta) cages in KI6 cells after 7.5 h of induced KLHL12 overexpression. (i) Overview of an area of the cell imaged by STORM. Small (80-100 nm diameter) canonical SEC31A puncta did not co-localize with PC1 (0%, n=76 puncta, N=8 experiment). Examples in dashed inset are magnified in (ii) as merged image, (iii) as SEC31A, and (iv) as PC1. Large SEC31A puncta of diameter above 300 nm colocalized with PC1 (76.3%, n=36 puncta, N=8 experiments). An example in solid inset is magnified in (v) as merged image, (vi) as SEC31A, (vii) as PC1. A virtual cross-section in the XZ plane of the large COPII vesicle shown in (viii) as merged image (ix) as SEC31A; (x) as PC1. (B) Triple-color 3D STORM imaging shows endogenous PC1 (cyan) encapsulated by endogenous KLHL12 (yellow) and SEC31A (magenta) in Saos-2 cells grown at steady state. XY maximum projection of the overview is shown in (i), and of a magnified vesicle was shown in (ii-v). Virtual cross-sections in the XZ plane were shown in (x-xii) and YZ in (vi-ix). Merged three-color channel is shown in (i, ii, vi, and x), SEC31A in (iii, vii, xi), PC1 in (iv, viii, xii), and KLHL12 in (v, ix, xii). Scale bars in (A i): 1 μm, (B i): 2 μm, and in magnified views: 100 nm

### PC1 containing giant COPII vesicles exhibit movement

The dynamic nature and organization of ER-to-Golgi transport have been visualized previously with the use of fluorescent labeling and time-lapse microscopy (Scales et al., 1997; Presley et al., 1997; Shima et al., 1999). We sought to understand whether the large COPII vesicles were capable of functioning as mobile transport carriers independent of the ER. The localization of large COPII vesicles relative to the ERES and ER was examined in KI6 cells by dual-color confocal and STORM microscopy using antibodies against SEC31A, SEC16A, a scaffolding protein at the ERES, and KDEL, the ER retrieval signal at the C-terminus of ER resident proteins. Images from both approaches showed clear separation of large COPII and ERES or ER marker proteins (Fig. 4A-B). Large puncta positive for both SEC31A and SEC16A were also observed by confocal microscopy (Fig. 4A). These coincidently labeled structures may represent pre-budding complexes of COPII vesicles that remained attached to the ER.

**Figure 4.**
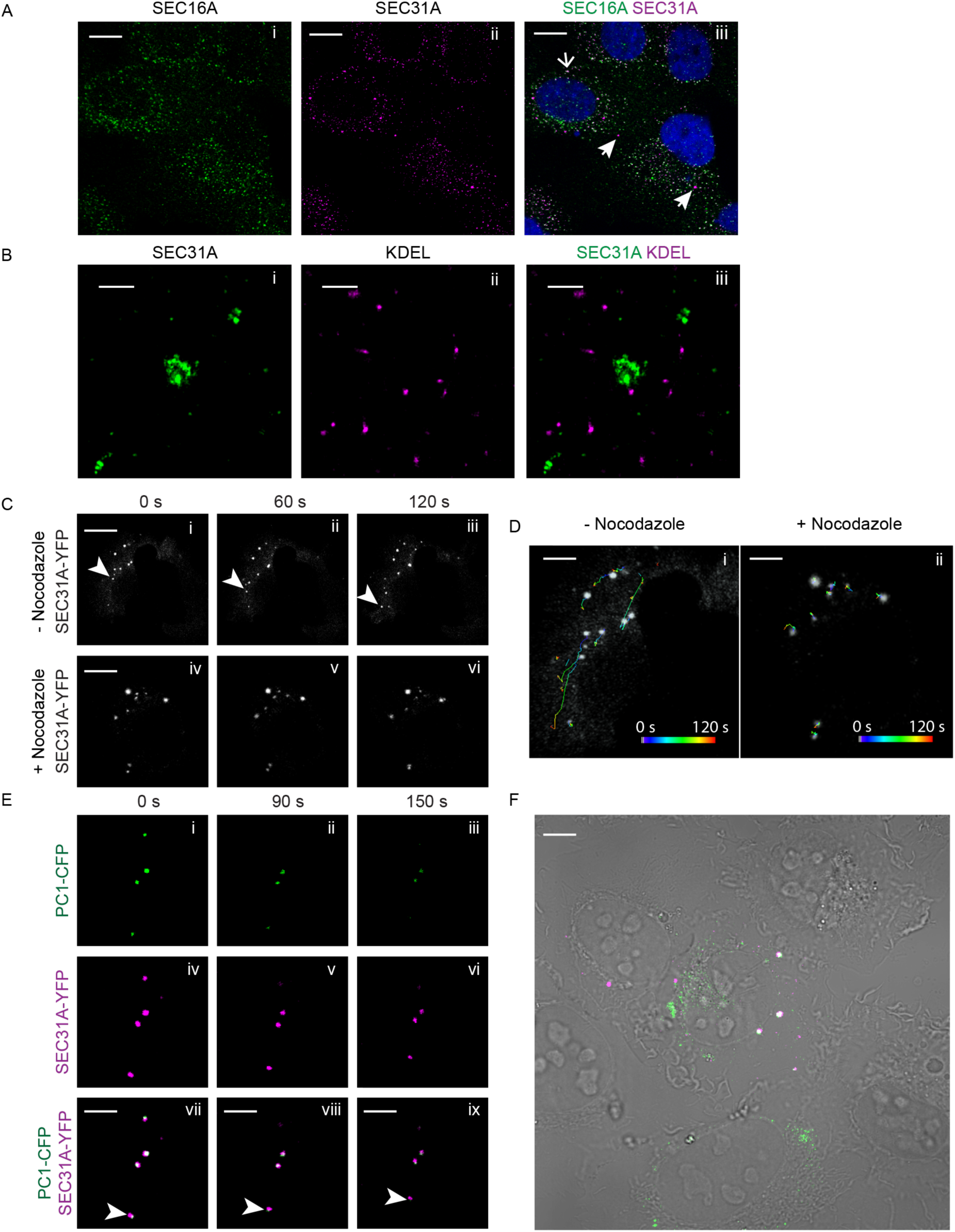
Procollagen carrying vesicles exhibit movement. (A) Confocal microscopy imaging on KI6 cells treated with doxycycline for 7.5 h were labeled with SEC16A and SEC31A antibodies. Open arrow in merged image points to a co-localized spot; closed arrow points to a large SEC31A-labeled vesicle not co-localizing with SEC16A. (B) Two-color STORM imaging was performed on KI6 cells induced for KLHL12 overexpression for 7.5 h and then labeled with KDEL and SEC31A antibodies. Panels show magnified XY view of (i) a SEC31A (green)- labeled large COPII cage (ii) KDEL marker labeling ER (magenta) and (iii) merged image of the two channels. (C) KI6 cells were transfected with YFP-tagged SEC31A and induced for KLHL12 expression for 7.5 h. (i-iii) show time lapse imaging of SEC31A-YFP puncta in cells not treated with nocodazole. Arrow indicates a puncta exhibiting long-range transport (>10 μm distance). n=3 (iv-vi) show time lapse images of SEC31A-YFP puncta after the treatment of 5 μΜ nocodazole for 30 min. (D) Images shown are the first frame of the time-lapse movie overlayed with the trajectories of SEC31A-YFP puncta in the absence (i) and presence (ii) of nocodazole. Trajectories in time (0-120 s) is represented by color according to the scale bar in (D). n=3 (E) Time lapse imaging of SEC31A-YFP and PC1-CFP positive puncta in KI6 cells. Arrows point to a vesicle that displays a spatial displacement in time. (F) Overview of the cell imaged in (E); overlay of merged fluorescent image over DIC image shows PC1-CFP in green and SEC31A-YFP in magenta. Scale bars in (A): 5 μm, (B): 500 nm, (C):10 μm, (D-F): 5 μm.

We then visualized the spatio-temporal dynamics of these vesicles by imaging YFP-tagged SEC31A labeled structures with time-lapse microscopy. KLHL12 was induced in KI6 cells for 7.5 h, which were then treated with ascorbate for 10 min to promote the formation of hydroxyproline residues essential to the folding and ER exit of PC1 (Stephens and Pepperkok, 2002). We observed large COPII vesicles exhibiting both long-range (> 2 um displacement) and short-range (< 2 um displacement) transport as previously observed for fluorescently-tagged small COPII vesicles (Fig. 4C-D, Video 2, Stephens et al., 2000). Long-range transport was not evident in cells treated with nocodazole, a microtubule-depolymerizing agent, as determined by particle tracking of individual vesicles (Fig. 4C-D, Video 3). The long-range mobility of large COPII vesicles was observed to be independent of a CFP-tagged ER marker (Stephens et al., 2000), which is consistent with our observations in fixed samples (Video 4). To confirm that these large mobile COPII vesicles are PC1 carriers, we tracked vesicles positive for both YFP-tagged SEC31A and CFP-tagged PC1, which also exhibited directional movements (Fig. 4E,Video 5).

### COPII is required to package PC1 into vesicles that bud from the ER in a cell-free reaction

To further investigate whether the large cargo PC1 exits the ER within COPII-coated vesicles, we modified a cell-free reaction designed to detect the formation of transport vesicles that bud from ER membranes in a preparation of permeabilized cells (Fig. 5A, Kim et al., 2005, Merte et al., 2010). Donor ER membrane was prepared from IMR-90 cultured human fibroblasts and incubated at 30°C with purified recombinant human COPII proteins, cytosol, and nucleotides for 1 h to allow the formation of transport vesicles. Budded vesicles were isolated by a modification of our previous procedures to allow the detection of large vesicular carriers. A lower centrifugal speed of 7000 xg was used to sediment donor membrane from a complete incubation, and the supernatant fraction, which contained budded vesicles, was applied to the bottom of an Optiprep step flotation gradient. After a high-speed centrifugation step of 250,000 xg, 12 fractions were collected and their contents were analyzed by immunoblotting (Fig. 5B). Because lipid vesicles are buoyant, they floated to the top of the gradient as shown by the enrichment of a standard COPII cargo SEC22B in fraction 1. In contrast, a soluble cytosolic protein marker, vinculin, was observed at the bottom of the gradient in fractions 7-12. Most PC1 signal floated to the top of the gradient and co-fractionated with SEC22B, indicating that most PC1 in the reaction was associated with membrane.

**Figure 5.**
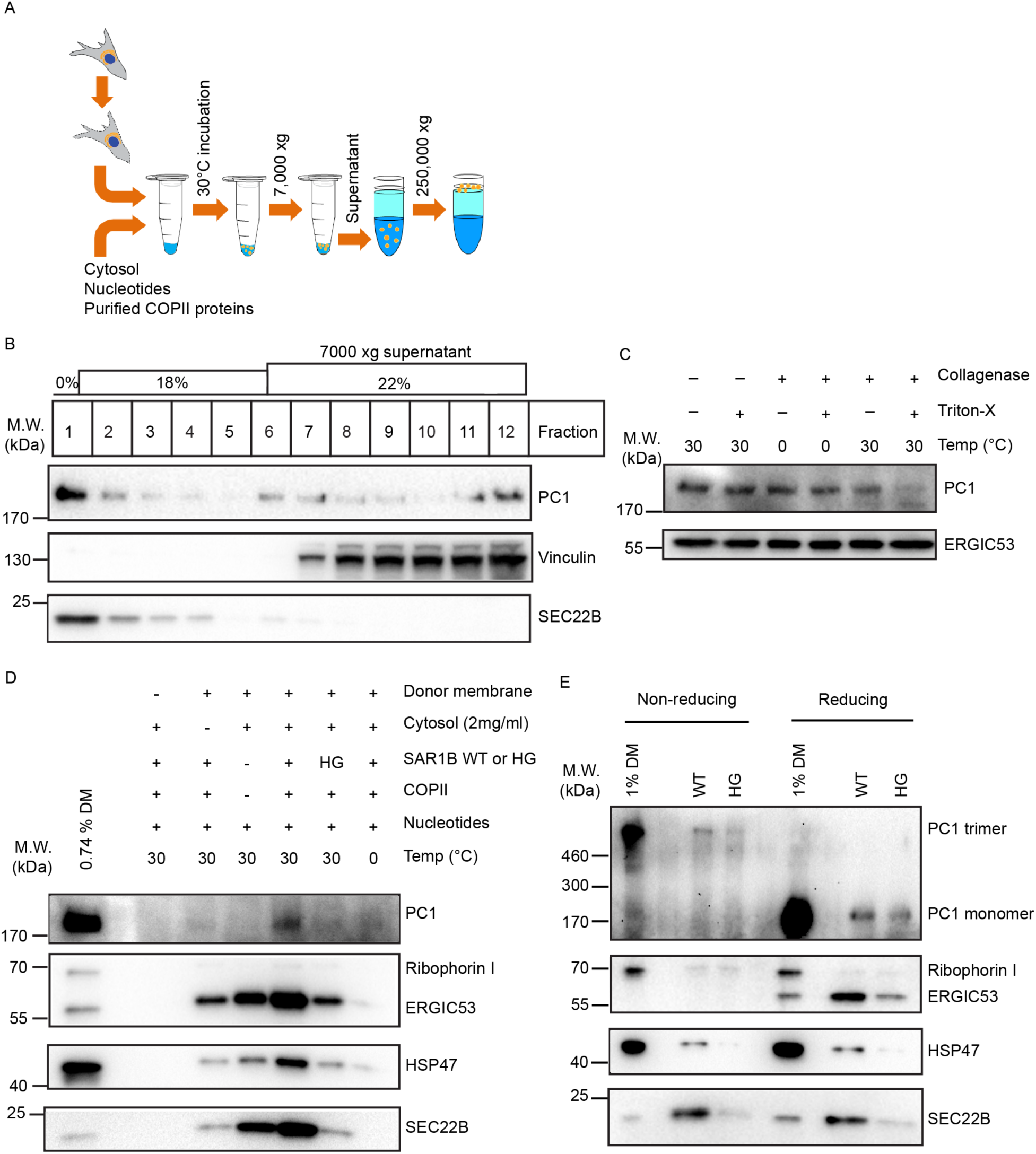
COPII is required to export PC1 in vesicles that bud from the ER in a cell-free reaction. (A) Scheme depicting the experimental procedure of the cell-free vesicle budding reactions. Briefly, donor membrane prepared from IMR-90 cells were incubated at 30°C with cytosol (from HT1080 or HTPC1.1 cells), nucleotides (an ATP regenerating system and GTP), and purified recombinant human COPII proteins. Vesicles in 7000 xg supernatant fractions from budding reactions were isolated by flotation. (B) Fractions (12) were taken from the top of an Optiprep gradient after flotation and analyzed by immunoblotting. SEC22B is found in conventional COPII vesicles and serves as a positive control. Vinculin serves as a control for cytosolic proteins that do not float with vesicles. n=3. (C) Top floated fraction was treated with or without collagenase (0.1 U/μl) in the presence or absence of the detergent Triton X (1%) at the indicated temperature for 10 min. ERGIC53 is found in conventional COPII vesicles and serves as loading control. n=4. (D) Budding requirements of PC1 and HSP47 were assessed under different incubation conditions. The top fraction after flotation was taken from each sample and analyzed by immunoblotting. Ribophorin I is an ER resident protein that serves as a negative control. ERGIC53 and SEC22B are found in conventional COPII vesicles and serve as positive controls. n=3. (E) To detect trimeric PC1, we treated donor membrane or floated fractions with non-reducing denaturing sample buffer before gel electrophoresis. Donor membrane or floated fractions treated with reducing denaturing sample buffers were used as controls. n=2

To test whether PC1 detected in the floated fractions was packaged inside of intact membrane vesicles, we employed a collagenase protection assay. When the top floated fraction was treated with collagenase at 30°C, most of the PC1 was protected from collagenase digestion and only became susceptible when detergent (1% Triton-X) was included to lyse the membrane (Fig. 5C). Because collagenase digestion was specific to collagens, another standard COPII cargo, ERGIC53, was used as loading control.

We tested the requirement for COPII in PC1 packaging by supplementing the reaction with the inhibitory SAR1B H79G GTPase-defective mutant protein in place of wild-type SAR1B (Fig. 5 D). Significantly less PC1 was detected in the top floated fraction, consistent with the observation that microinjection of SAR1 H79G into live cells arrested PC in the ER (Stephens and Pepperkok, 2002). As further controls, the cell-free reaction was carried out in the absence of donor membrane, cytosol, or purified recombinant COPII proteins, respectively. Immunoblotting of the top floated fractions showed that donor membrane, cytosol, purified COPII proteins and a physiological temperature of 30°C were required to reconstitute export of PC1 (Fig. 5D). PC1 packaging appears to be completely dependent on both cytosolic proteins and COPII for packaging; thus, soluble factor(s) in addition to the COPII coat may be essential to sort PC1 for transport.

PC1 is known to trimerize in the ER, and only the correctly folded PC1 trimers are chaperoned by HSP47 to exit the ER *en route* to the Golgi membrane. Consistent with our observation in cells where HSP47 was found to be present on the large COPII vesicles, the budding of HSP47 was also observed in the cell-free reaction, which required similar conditions as the budding of PC1 (Fig. 5D, Fig. 1C). To further demonstrate the physiological relevance of our cell-free reaction, we analyzed the content of the floated fractions under a non-reducing, denaturing condition, which would preserve the inter-chain disulfide bonds and allow the detection of trimeric PC1 (Fig. S2A). In cell-free reactions supplemented with wild-type SAR1B, lower mobility PC1 ~ 570 kDa (the predicted size of PC1 trimer) was detected in the floated fraction under the non-reducing, denaturing condition, whereas monomeric PC1 ~190 kDa was detected under a reducing, denaturing condition (Fig. 5E). Consistent with observations under the reducing, denaturing condition, the amount of trimeric PC1 observed in the floated fraction was significantly decreased on incubation with SAR1B H79G (Fig. 5E). Thus, we conclude that the COPII-dependent packaging of trimeric PC1 and its chaperone HSP47 is sustained in our cell-free reaction.

### PC1 is exported out of the ER in large COPII-coated vesicles

To visualize the morphology of vesicular PC1 carriers in the floated fraction, we modified the cell-free reaction so that both the COPII coat and the large cargo PC1 were fluorescently labeled. For fluorescence visualization of isolated COPII-coated vesicles, we added purified Alexa Fluor 647-conjugated SEC23A/24D to the reaction mixture (Bacia et al., 2011). For fluorescence visualization of PC1, we prepared donor membrane from cells that were transiently transfected with a C-terminally GFP-tagged construct of the pro-α1(I) chain of human PC1. This construct was shown to produce GFP signal that exits the ER in cells incubated in ascorbate-containing medium (Stephens and Pepperkok, 2002). We confirmed that the GFP tag did not interfere with trimerization and ER to Golgi transport of PC1-GFP, as GFP signal was found at the Golgi apparatus after the addition of ascorbate (Fig. S4A). Moreover, PC1-GFP was packaged in a COPII-dependent manner in the vesicle budding reaction as detected by immunoblotting of GFP in the floated fraction (Fig. S4B).

Vesicles observed by structured illumination microscopy (SIM) displayed PC1 co-localized with large COPII-coated structures of around 400 nm in diameter (Fig. 6A). With the resolution of SIM, the PC1-GFP signal appeared fully within the signal of SEC23A/24D. Smaller COPII structures devoid of PC1-GFP, likely to be conventional COPII vesicles, were also observed. Not all large COPII structures observed contained the PC1-GFP signal (quantified in Fig. 6Biii, iv), possibly because only 10-30% of the donor membrane contained GFP signal due to low transfection efficiency. In addition, PC1 may not be the only large cargo exiting the ER in large COPII-coated vesicles. Also, not all PC1-GFP observed co-localized with SEC23A/24D. This non-overlapping GFP signal may represent PC1 not enclosed within membranes, possibly coinciding with the amount of PC1 that was sensitive to collagenase digestion (Fig. 5C); alternatively, the signal may emanate from transport vesicles from which the COPII coat had been shed (Fig. 3B, Fig. S3, Fig. S5A).

**Figure 6.**
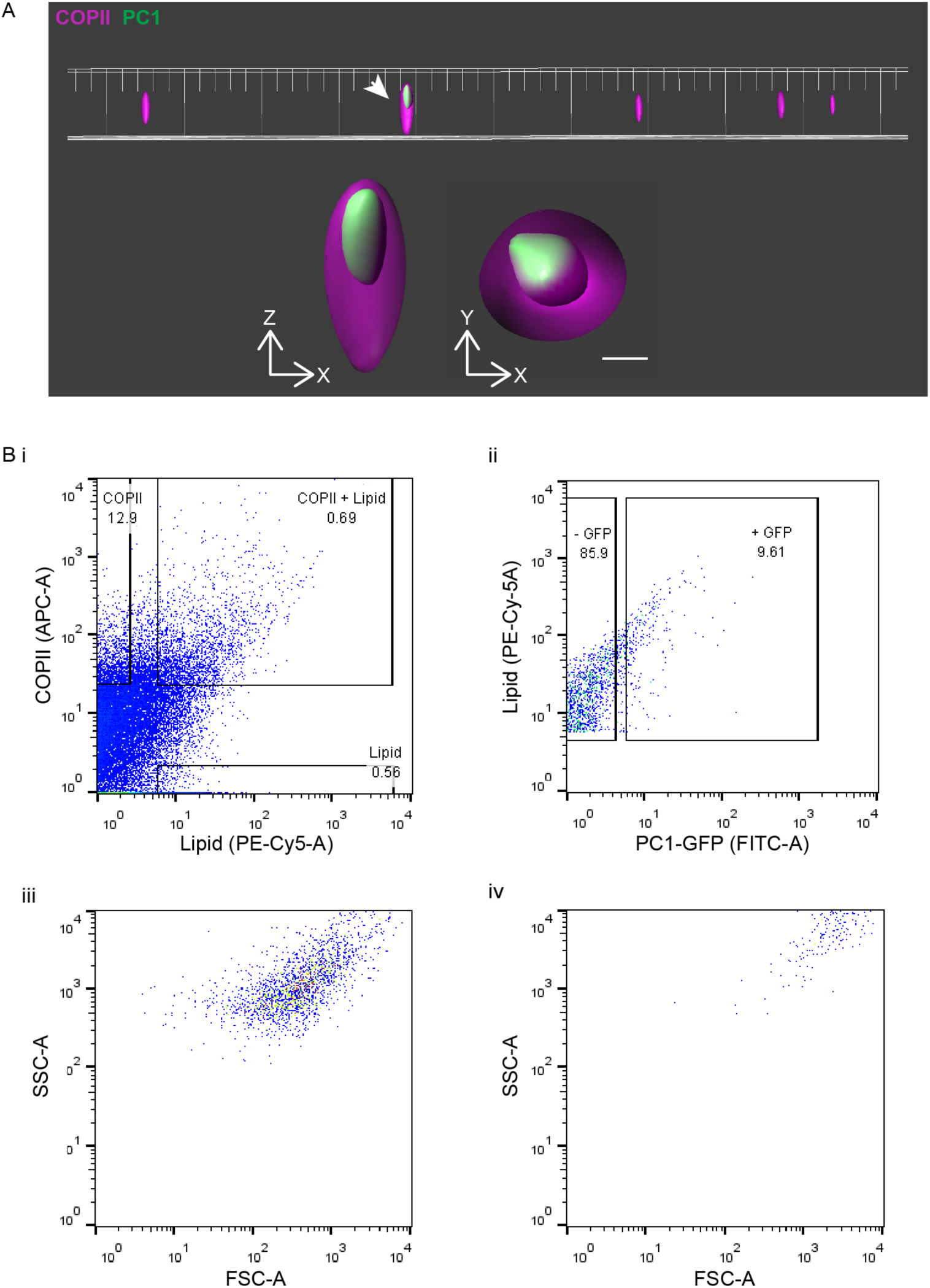
PC1 is exported out of the ER in large COPII-coated vesicles. (A) A representative field of COPII vesicles isolated from cell-free vesicle budding reaction visualized by SIM. One of them indicated by the arrow contained PC1-GFP (green) that was entirely encapsulated by the COPII inner coat proteins SEC23A/24D-647 (magenta). The magnified XZ and YZ views are displayed underneath the overview. Around 20 images were collected from each experiment with fluorescence controls. n=3. Bar: 200 nm. (B) Relative size of COPII carriers of PC1 compared to regular COPII vesicles quantified by flow cytometry, and numbers in graphs represent the percentage of particles in the respective subpopulation. (i) Singlets were plotted for the fluorescence intensities of the COPII coat proteins SEC23A/24D-647 (APC-A) and the lipid dye FM4-64 (PE-Cy5-A). (ii) Particles positive for both “COPII” and “lipid” are COPII vesicles and were plotted for the fluorescence intensity of PC1-GFP (FITC-A). (iii-iv) The values of side scatter (SSC-A) versus forward scatter (FSC-A) of COPII vesicles that are PC1-GFP negative (iii) or GFP positive (iv) are plotted. n=3.

We developed an unbiased flow cytometry approach to quantify the relative fluorescence signal associated with both conventional and large COPII vesicles. Single particles (singlets) were gated for their content of fluorescently labeled COPII proteins SEC23A/24D and the fluorescent lipid dye FM4-64. Particles that satisfied such criteria accounted for about 0.7% of all singlets detected in the 7000 xg supernatant fraction (Fig. 6Bi). Of these COPII-coated vesicles, 9.6% were GFP positive indicating that they carried the large cargo PC1-GFP (Fig. 6Bii). This population of collagen carrying COPII vesicles exhibited higher side and forward light scatter, both of which positively correlate with larger particle sizes (Fig. 6Biii-iv). The sizes of reconstituted COPII-coated vesicles were estimated using Nanoparticle Tracking Analysis (NTA), which tracks and analyzes the Brownian motion of each particle (Fig. S5A, Dragovic et al., 2011). COPII-coated vesicles in the range of 50-150 nm were ~17.6 x more prevalent than larger COPII-coated vesicles in the range of 300-1000 nm. A medium sized category of 150-300 nm was also observed and it was ~4.3 x less prevalent than the 50-150 nm category and 4.1 x more in number than the larger vesicles. The existence of medium- to large-coated membrane vesicles with a coat of 10-15 nm was confirmed independently by thin-section EM (Fig S5B)

### Recruitment of functional KLHL12 to COPII-coated vesicles

To test whether our cell-free COPII vesicle formation reaction reconstitutes the cellular behavior of KLHL12, we supplemented the reaction with cytosol collected from cells transfected with either FLAG-tagged wild-type KLHL12 or a mutant, FG289AA, which fails to promote the formation of large COPII vesicles (Jin et al, 2012) (Fig. 7). Densitometry analyses of immunoblots showed 6.8 x more wild-type KLHL12 in the vesicle floated fraction compared to the FG289AA mutant protein (p<0.0001, paired t-test, n=7) (Fig. 7A,B). This result coincided with our previous observation that KLHL12 FG289AA mutant protein rarely co-localized with COPII markers in cells (Jin et al., 2012). The relative budding efficiency of PC1 and control COPII cargos were also calculated from densitometry analyses of immunoblots, and PC1 budding was only marginally stimulated by KLHL12 wild-type cytosol compared to KLHL12 FG289AA cytosol (p=0.0412, paired t-test, n=7) (Fig. 7A,C). Thus, although we have reconstituted the association of functional KLHL12 with budded vesicles, we were unable to assess the role of this protein in the capture of PC1 in our cell-free reaction. There may be other rate-limiting components, such as the calcium-binding co-adaptors PEF1 and ALG2, which may fail to be recruited to the site of vesicle budding under the conditions of this incubation (McGourty et al., 2016).

**Figure 7.**
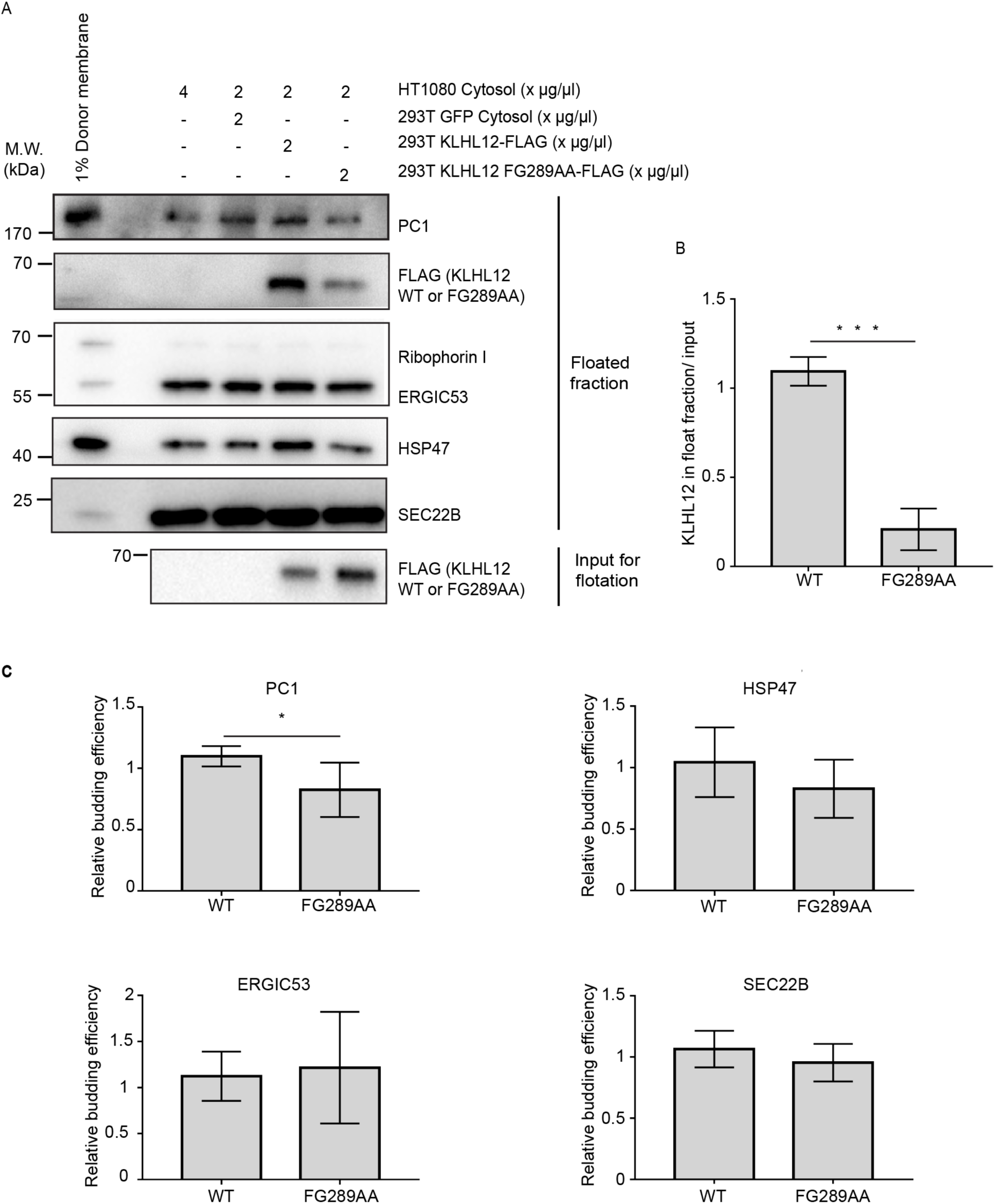
The cellular behavior of KLHL12 is recapitulated in the cell-free reaction. (A) Budding reactions were supplemented with 2μg/ul 293T cytosol containing FLAG-tagged KLHL12 wild-type or FG289AA mutant in addition to 2μg/u1 HT1080 cytosol. Cytosol (293T cells) that contained GFP and 4μg/μ1 HT1080 cytosol were used as controls with endogenous levels of wild-type KLHL12. Top fractions were collected after density gradient flotation for immunoblotting analyses. A representative experiment out of 7 independent repeats. (B) Relative enrichment of KLHL12 detected in the floated fraction. Densitometry quantification of FLAG intensities in the floated fraction was divided by the corresponding FLAG intensity in the 7000 xg supernatant (input for flotation). Error bars: SD. Paired t-test. p<0.0001. n=7. Average of wild-type Ι average of FG289AA = 6.80. (C) Relative budding efficiency of PC1, HSP47, ERGIC53 and SEC22B in reactions that were supplemented with 293T cytosol that contained wild-type KLHL12 or KLHL12 FG289AA. Vesicle budding efficiencies were calculated from densitometry quantification relative to the control lane where 293T cytosol that contained GFP was supplemented. Paired t-test was used to analyze the difference between wild-type and FG289AA samples of each cargo and the p values were 0.0412 for PC1, 0.1215 for HSP47, 0.5307 for ERGIC53, and 0.2993 for SEC22B. Error bars: SD. n =7.

## Discussion

The function of canonical (60-80 nm) COPII vesicles in conventional cargo transport is well established: COPII subunits concentrate cargo proteins into vesicles that bud from the ER and form a structure which dictates the size and contour of the bud and completed vesicle (Zanetti et al., 2012). Although a wide range of studies has shown that PC secretion is COPII dependent, it has remained unclear whether COPII coats the membranes surrounding the large cargo or instead plays an indirect role in cargo packaging and transport carrier biogenesis. A primary requirement for the transport of large cargos would be to overcome the structural constraints of a small cage (Noble et al., 2013). KLHL12 was previously shown to fulfill this requirement, as overexpressing KLHL12 induced the formation of large COPII structures (Jin et al., 2012). Here, we confirmed that these large COPII-coated structures appeared hollow, presumably with a space to contain large cargos (Fig. 2A-B). Similar observations were also made in the PC1-secreting osteosarcoma cells Saos-2 without any overexpression of KLHL12 (Fig. 2C, Fig. 3B). Notably, CLEM analysis of these structures in Saos-2 showed a COPII-coated single membrane bilayer envelope surrounding a lumenal space filled with PC1 (Fig. 2C).

Evidence for the contrary model that PC is transported out of the ER in non-COPII carriers came from studies of fibroblasts released from a PC1 hydroxylation block (Mironov et al., 2003). This study reported that PC1 exits from saccular extensions of the ER that appeared devoid of the COPII coat based on immunofluorescence microscopy and immuno-gold EM studies. These conclusions were based on the use of a polyclonal antibody, LF-68, which was generated against synthetic peptides of a unique sequence selected from the C-telopeptide (non-triple helical segment exposed on the C-terminus of mature collagen after a C-terminal propeptide is cleaved) of human α1(I) collagen (Fig. S2A, Fisher et al., 1995; Bernstein et al., 1996). Here, using two monoclonal antibodies, QED-42024 and sp1.d8, directed against C- and N-terminal propeptides cleaved from triple helical PC1 respectively (Fig. S2A, Personal communication, QED Biosciences;Foellmer et al., 1983), we observed co-localization with large COPII structures in KI6 cells where both PC1 and KLHL12 were overexpressed (Fig. 1A-C, Fig. 3A), in natural collagen I secreting svIMR-90 cells that overexpressed KLHL12 (Fig. S2C), and in natural collagen I secreting Saos-2 and IMR-90 cells where PC1 and KLHL12 are expressed endogenously (Fig. 2C, Fig. 3B, Fig. S2D). We have independently confirmed the results of Mironov et al. (2003) using the same polyclonal antibody (LF-68) for IF localization (Fig. S2D). However, we suspect the C-telopeptides are not fully accessible in non-denatured PC1 trimers, possibly hindered by the presence of C-terminal propeptides, which are cleaved in post-Golgi vesicles or after secretion to the extracellular space (Fig. S2A). Thus this polyclonal antibody may have limited utility in the immunofluorescence detection of folded PC1 species in early secretory transport vesicles.

Independent of PC1 antibody labeling, the co-localization of PC1-CFP and the COPII coat protein SEC31A-YFP was also observed in live cells (Fig. 4E). Moreover, the PC-specific chaperone HSP47 also co-localized with KLHL12 and COPII (Fig. 1C). HSP47 binds to trimerized PC in the ER and assists its correct folding by preventing lateral aggregation (Nagai et al., 2000; Tasab et al., 2000; Ono et al, 2012; Widmer et al., 2012). This chaperone traverses the early secretory pathway with PC to ERGIC or cis-Golgi membranes, where it is released due to lower pH, before being recycled back to the ER via its C-terminal RDEL sequence (Satoh et al., 1996, Oecal et al., 2016). In HSP47 knockout mouse embryonic fibroblasts, mis-folded PC1 trimers were observed to accumulate in the ER and were degraded by autophagy, independent of the ERAD-mediated clearance of monomeric PC1 aggregates (Ishida et al., 2006). Hence, the observation of HSP47 co-localized with large COPII confirms that these large COPII structures contain correctly folded triple helical PC1 and are physiologically relevant.

STORM and CLEM analyses revealed the ultrastructural details of these large COPII structures: collagen became completely encapsulated in a large COPII coat during vesicle formation, suggesting that COPII proteins acted directly by forming a complete cage around a bilayer vesicle that carried PC1 (Fig. 2). Furthermore, fluorescently labeled large COPII vesicles showed microtubule-dependent vectorial movement in live cells (Fig. 4C-E). To sum up, *bona fide* large COPII-coated membrane bilayer carriers of PC1 were observed in fixed as well as in live cells.

In an independent effort to test the role of COPII-coated vesicles in the sorting of PC1, we employed a cell-free transport vesicle budding assay that was previously developed in our lab to probe the requirements for sorting of conventional COPII cargo proteins (Kim et al., 2005; Merte et al., 2010). Here, we report an alternative fractionation method to detect the ER export of the large cargo PC1 and its chaperone HSP47 (Fig. 5). Using the combination of a buoyant density flotation protocol, a collagenase protection assay, and SIM, we showed that PC1 exits the ER inside of large membrane vesicles that are coated with COPII proteins (Fig. 5B-C, Fig. 6A). Furthermore, the packaging of PC1 requires the addition of recombinant COPII proteins, as well as of other unspecified cytosolic protein(s) and depends upon the GTPase activity of SAR1, the GTP-binding protein that initiates the assembly of the COPII coat (Fig. 5D).

Although we were unable to identify a function for KLHL12 in our cell-free reaction, we observed the wild-type but not an ubiquitylation-defective mutant form of the protein recruited to budded vesicles (Fig. 7). The requirement for cytosolic proteins in addition to COPII is consistent with reports in the literature of other factors such as Sedlin, Sly1, TFG, ALG2 and PEF1, in the export of PC from the ER (Veditti et al., 2012; Noguiera et al., 2014; McCaughey et al., 2016, McGourty et al., 2016). Further resolution of the cytosolic proteins required in the cell-free reaction may illuminate those that cooperate with KLHL12 to stimulate PC1 packaging.

In summary, we examined the role of COPII during the ER export of large cargo PC1 using a combination of *in vivo* morphological analyses and *in vitro* reconstitution studies. Our results support the conventional model where COPII participates directly by forming a large COPII-coated membrane vesicle to transport bulky cargo out of the ER. Although the exact mechanism by which the size of COPII vesicles is regulated awaits further investigation, the cell-free PC1 budding reaction described here should aid in such studies.

## Materials and Methods

### Antibodies and plasmids

Commercially available antibodies used for immunofluorescence (IF) and immunoblotting (IB) were as follows: mouse anti-PC1 (QED Biosciences, San Diego, CA, USA clone # 42024) 1:200 (1mg/ml stock concentration) for IF, rabbit anti-SEC31A (Bethyl Laboratories, Montgomery, TX, USA for IF at 1:200 for confocal and 1:2000 for STORM), mouse anti FLAG (Thermo Fisher Scientific, Waltham, MA, USA, 1:5000 for IB), goat anti-FLAG (Novus Biologicals, Littleton, CO, USA for IF at 1:1000 for confocal and 1:5000 for STORM), chicken anti-KLHL12 (Novus Biologicals, for IF at 1:200 for confocal and 1:2000 for STORM), mouse anti- KLHL12 (Cell Signaling Technology, Danvers, MA, USA clone#2G2, 1:1000 for IB), rabbit anti-calnexin (Abcam, Cambridge, MA, USA 1:200 for IF), mouse anti-HSP47 (Enzo Life Sciences, Farmingdale, NY, USA 1:200 for IF; 1:5000 for IB), sheep anti-TGN46 (AbD Serotec, Raleigh, NC, USA, 1:200 for IF), mouse anti-KDEL (Enzo Life Sciences, clone 10C3, for IF at 1:200), mouse anti-Vinculin (Abcam, 1:5000 for IB) and rabbit anti-GFP (Torrey Pines Biolabs, Secaucus, NJ, USA 1:1000 for IB). Rabbit anti-ribophorin I, ERGIC53, SEC22B were made in house and used at 1:5000 for IB. Rabbit anti-PC1 LF-41, 67, 68 antibodies were obtained as a gift from Larry Fisher (Fisher et al., 1989; Fisher et al., 1995; Bernstein et al., 1996) at NIH/NIDCR. LF-67 and LF-68 were raised against the same synthetic peptide of the human α1(I) collagen C-telopeptide (aa 1192-1218) (Bernstein et al., 1996) and LF-68 was used at 1:1000 for IF only to repeat previous reports (Mironov et al., 2003; Jin et al., 2012). LF-41 was raised against a synthetic peptide of the C-terminus of the human α1(I) collagen (aa 1443-1464) (Fisher et al., 1989), and it was used for all anti-PC1 IB at 1:5000, but not for IF in this report. The mouse anti-PC1 antibody sp1.d8 was purified in house from culture medium of the mouse hybridoma cells obtained from Developmental Studies Hybridoma Bank at the University of Iowa using standard procedures and used at 3.75 ng/ul for IF. Expression constructs for SEC31A-YFP, PC-CFP, PC-GFP and pCFP-ER (encodes a fusion protein consisting of enhanced cyan fluorescent protein (CFP) flanked by a N-terminal signal peptide of calreticulin and a C-terminal ER retrieval sequence, KDEL) were kindly provided by David Stephens lab (University of Bristol, UK).

### Cell culture, transfection, and drug treatments

Human lung fibroblasts IMR-90 and svIMR-90, IMR-90 immortalized with SV-40, were obtained from Coriell Cell Repositories at the National Institute on Aging, Coriell Institute for Medical Research (Camden, New Jersey, USA). Human osteosarcoma Saos-2 and U-2OS, and human fibroscarcoma HT-1080 were obtained from ATCC (Manassas, Virginia, USA). IMR-90, svIMR-90, Saos-2, U-2OS and HT-1080 were maintained in DMEM plus 10% FBS. The HT-PC1.1 cells line was generated from HT-1080 as previously described in Jin et al., 2012 by stable expression of COL1A1 in a pRMc/CMV plasmid (gift from Neil Bulleid lab, University of Glasgow, UK) and maintained in 0.4mg/ml G418. The doxycycline-incudcible KLHL12- 3xFLAG stable cell line (KI6) was generated through sequential clonal selection of HT-PC1.1 cells that stably integrated pcDNA6/TR and KLHL12-3xFLAG in a pcDNA5/FRT/TO vector (Flp-In T-REx Core Kit, Thermo Fisher) which were selected in the presence of 6μg/ml blasticidin and 0.2mg/ml hygromycin, respectively. Cells were kept in 37°C incubator with 5% CO2. Transfection of DNA constructs into svIMR-90 and KI6 cells was performed using lipofectamine 2000 as detailed in the manual provided by Invitrogen (Carlsbad, CA, USA). Transfection of DNA constructs to Saos-2, svIMR-90, and U-2OS cells for donor membrane preparation and HEK293T cells for cytosol preparation was performed using polyethylenimine (PEI) at a DNA to PEI ratio of 1:3. Ascorbate treatment used 0.25 mM ascorbic acid (Sigma-Aldrich, St. Louis, MO, USA) and 1 mM ascorbic-2-phosphate (Sigma-Aldrich) were used for ascorbic treatment. Doxycycline (Sigma-Aldrich) was used at 1 μg/ml. BFA (Sigma-Aldrich) was used at 10 μg/ml. Nocodazole was used at 5 μM (Sigma-Aldrich).

### Immunofluorescence

Cells growing on poly-lysine coated glass coverslips were fixed in 4% PFA for 20 min at RT or 30 min at 4°C, washed 5 times with PBS and then incubated with permeabilization buffer (PBS containing 0.1% TritonX-100 and 0.2M glycine) at RT for 15 min. Cells were then incubated with blocking buffer (0.5% BSA in PBS) for 30 min at RT followed by incubation for 1h each at RT with primary antibody and secondary antibody subsequently. Antibody incubations were followed by 5 washes with PBS. Coverslips were mounted in ProLong-Gold antifade mountant with DAPI (ThermoFisher Scientific) overnight, before imaging. Images were acquired using Zen 2010 Software on Carl Zeiss LSM 710 confocal microscope system. The objectives used were Plan -Apochromat 100X, 1.4 NA and Plan-Apochromat 100X, 1.4 NA. The excitation lines and laser power used were 488 (4%), 543 (6%), 405 (2%) and 633 (6%). Co-localization analysis was performed using the “Spots” detection algorithm in the image-processing module, Imaris 8.1.2. Briefly, a total number of KLHL12 labeled puncta in the Z stack was obtained and the percentage of puncta positive for both KLHL12 and PC1 was calculated.

### Immunoblotting

Standard immunoblotting procedures were followed. Briefly, samples were heated at 65°C for 10min, resolved on 4-20% polyacrylamide gels (15 well Invitrogen; 26 well Bio-Rad), and transferred to PVDF (EMD Millipore, Billerica, MA, USA). The PVDF membrane was incubated with antibodies and bound antibodies were visualized by the enhanced chemiluminescence method (Thermo Fisher Scientific) on a ChemiDoc Imaging System (Bio-Rad, Hercules, CA, USA) with ImageLab software v4.0 (Bio-Rad).

### Immunoelectron microscopy

KI6 cells were grown on 35 mm glass bottom dishes (MatTek Corporation, Ashland, USA), fixed with 4% PFA and 0.05% glutaraldehyde for 30 min. The cells were washed with PBS and incubated with blocking buffer (0.5% BSA, 0.02% saponin in PBS) at RT for 20 min. Primary antibody labeling was done at RT for 1h followed by overnight incubation at 4°C in the presence of saponin. Secondary antibody labeling (1: 50 dilution of 1.4 nm Nanogold conjugated Goat anti-mouse Fab’ fragment, Nanoprobes Inc.) was done at RT for 2h. The Nanogold particles were gold enhanced in dark for 5-7 min per manufacturer’s instructions using GoldEnhance™ (Nanoprobes, Yaphank, NY). Cells were post-fixed with 1% osmium tetroxide and processed for thin section EM.

### Correlative Light and Electron Microscopy (CLEM)

Saos-2 cells were grown on gridded 35 mm glass bottom dishes (MatTek Corporation, Ashland, USA) and fixed with PFA (4% in culture medium, Electron Microscopy Sciences, Hartfield, PA, USA). The cells were processed for immunofluorescence as above with the exception that 0.02% saponin was used as the permeabilizing agent, a concentration well below the established protocol which typically employs 0.1% saponin (Mironov et al., 2005). The cells displaying colocalized PC1/SEC31A spots were imaged on a Zeiss 710 Confocal Microscope using a C-Plan Apochromat 63X/1.2 W objective followed by marking the alphanumeric location using a 5X / 0.12 NA objective. A Z-stack series of the cell of interest was also collected. The cells were then fixed for 30 min in 0.1M cacodylate buffer, pH 7.2, containing 2% glutaraldehyde, and washed with 0.1 M sodium cacodylate buffer prior to post-fixation with 1% osmium tetroxide for 30 min on ice. This was followed by staining with 1 % aqueous uranyl acetate for 30 min at RT. For dehydration with progressive lowering of temperature, each incubation period was 10 min, with exposure to 35% ethanol at 4°C, to 50% ethanol and 70% ethanol at -20°C, and 95%, and 100% ethanol at -35°C. Cells were restored to RT in 100% ethanol before embedding in an Epon resin. The cell of interest was identified by the grid location on the resin and thin (70-100nm) serial sections were collected on Formvar-coated 200-mesh copper grids and post-stained with 2% aqueous uranyl acetate and 2% tannic acid. The sections were imaged at 120 kV using a Tecnai 12 Transmission Electron Microscope (FEI, Eindhoven, Netherlands). Regions of interest were identified manually by correlating the Z slices of the confocal stack with the serial sections and overlaying the fluorescent image over the TEM image by marking down prominent cell landmarks such as the nuclear boundary, vacuoles and mitochondria. The overlay was confirmed using the Icy software package (Chaumont et al., 2012).

### STORM imaging

Immunofluorescently labeled cell samples were mounted on glass slides with a standard STORM imaging buffer consisting of 5% (w/v) glucose, 100 mM cysteamine, 0.8 mg/ml glucose oxidase, and 40 μg/ml catalase in Tris-HCL (pH 7.5) (Rust *et al.* 2006; Huang *et al.* 2008), and sealed using Cytoseal 60 (Thermo Fisher Scientific). STORM imaging was performed on a home-built setup based on a modified Nikon Eclipse Ti-U inverted fluorescence microscope using a Nikon CFI Plan Apo λ 100x oil immersion objective (NA 1.45). Dye molecules were photo switched to the dark state and imaged using either 647- or 560-nm lasers (MPB Communications, Pointe-Claire, QC, Canada); the lasers were passed through an acousto-optic tunable filter and introduced through an optical fiber into the back focal plane of the microscope and onto the sample at intensities of ~2 kW cm^-2^. A translation stage was used to shift the laser beams towards the edge of the objective so that light reached the sample at incident angles slightly smaller than the critical angle of the glass-water interface. A 405-nm laser was used concurrently with either the 647- or 560-nm lasers to reactivate fluorophores into the emitting state. The power of the 405-nm laser (typical range 0-1 W cm^-2^) was adjusted during image acquisition so that at any given instant, only a small, optically resolvable fraction of fluorophore in the sample was in the emitting state. For 3D STORM imaging, a cylindrical lens was inserted into the imaging path so that images of single molecules were elongated in opposite directions for molecules on the proximal and distal sides of the focal plane (Huang et al, 2008). The raw STORM data was analyzed according to previously described methods (Huang et al, 2008; Rust et al, 2006). Data was collected at a frame rate of 110 Hz using an Andor iXon Ultra 897 EM-CCD camera, for a total of ~80,000 frames per image. Three-color imaging was performed on targets labeled by Alexa Fluor 647, CF680, and CF568 via sequential imaging with 647-nm and 560-nm excitation. With 647-nm excitation, a ratiometric detection scheme (Bossi *et al.* 2008; Testa *et al.* 2010) was first employed to concurrently collect the emission of single Alexa Fluor 647 and CF680 molecules. Emission of single molecules was split into two light paths (channels) using a long pass dichroic mirror (T685lpxr; Chroma, Bellows Falls, VT), each of which were projected onto one-half of an Andor iXon Ultra 897 EM-CCD camera. We performed fluorophore assignment by localizing and recording the intensity of each single molecule in the two channels. Excitation at 560 nm was subsequently used to image CF568 through the reflected light path of the dichroic mirror.

### Live cell Imaging, particle tracking and image analysis

KI6 cells were transfected with plasmids encoding SEC31A-YFP and PC1-CFP for a period of 9 h and induced for 7.5 h with doxycycline within that time period to induce the over expression of KLHL12. The cells were imaged using a temperature controlled Zeiss LSM 710 Axio Observer microscope (Plan-Apochromat 63X, 1.4 NA Oil DIC M27; standard Zeiss PMT detectors) at 37°C in pre-warmed Phenol Red-free culture medium. Main Beam splitters (MBS) were MBS-405 and MBS-458/514. Emission wavelengths collected were at 454-516 nm and 519-621 nm. Cells were imaged every 5 s for 2 min started at the end of 10 min treatment with ascorbate. For experiments requiring nocodazole treatment, the drug was added for 30 min before imaging.

Particle detection and tracking were performed using Imaris v8.1 (Bitplane USA, Concord, MA, USA) software. Images were first subjected to background subtraction and the “Spots” module was used to automatically detect point-like particles with spot diameter of 200 nm. Appropriate threshold values were confirmed. Object tracing through sequential time frames was done using an autoregressive motion particle-tracking algorithm. A maximum search distance of 5 μm was defined. A gap-closing algorithm was also implemented to link track segment ends to track segment starts. Track outputs were then visually inspected.

### Vesicle budding reaction

Vesicle budding reactions were performed as previously described in Kim et al., 2005 with the following modifications. Donor ER membrane was prepared fresh for each reaction by permeabilizing young (under PDL 37.5) IMR-90 cells that were grown (95% confluent in 3x10 cm dishes) were treated with 20 μg/ml digitonin (5 min on ice) in B88 (20 mM HEPES pH7.2, 250 mM sorbitol, 150 mM potassium acetate, 5 mM magnesium acetate) and washed with 0.5 M LiCl in B88 then B88 and resuspended in 200 μl B88-0 (20 mM HEPES pH7.2, 250 mM sorbitol, 150 mM potassium acetate) to a final concentration of OD_600_ between 2 to 6. Each 100 μl reaction contained ATP regeneration system (1 mM ATP, 40 mM creatine phosphate, 0.2 mg/ml creatine phosphokinase), 3 mM GTP, purified human COPII proteins (2 μg SAR1B, 1 μg SEC23A/24D, and 1 μg SEC13/31A), cytosol (2 μg/ul unless otherwise stated, see “cytosol preparation” for details), and a final concentration of 5 OD_600_/ml donor ER membrane in B88-0. Vesicles generated in vitro were isolated from the reaction mixture in 2 steps. First, donor membranes were sedimented by centrifuging twice at 7000 xg at 4°C for 5 min each in a swinging bucket rotor (Eppendorf S-24-11-AT). Then, 85 μl of the supernatant was mixed with 50 μl 60% OptiPrep (Sigma-Aldrich) gently until homogenous, placed at the bottom of a 7 x 20 mm tube (Beckman Coulter, Brea, CA, USA) and overlaid with 100 μl 18% and 10 μl 0% OptiPrep in B88. The OptiPrep gradient was centrifuged at 250,000 xg for 90min at 4°C (Beckman TLS-55 with adaptors for 7 x 20 mm tubes) with slow acceleration and deceleration, after which 20 μl fractions were collected from the top and mixed with sample buffer for immunoblotting analysis. When a vesicle budding reaction was analyzed by SIM or flow cytometry, the donor ER membrane was prepared from cells (2x10 cm 80% confluent Saos-2, 40% confluent U-2OS or 60% confluent svIMR-90 at the time of transfection) transiently transfected with 7.5 μg COL1A1-GFP per 10cm plate using PEI (Sigma-Aldrich) for 72 h when cells are 100% confluent. The reaction was supplemented with fluorescently labeled SEC23A/24D. Fluorescent SEC23A/24D was produced as previously described in Bacia et al., 2011 using Alexa Fluor 647 C2 Maleimide (Thermo Fischer Scientific).

### Protein purification

Human SAR1 proteins were overexpressed in *E. coli* and purified as cleaved GST-fusions, as described for hamster Sar1 purifications in Kim et al., 2005. Human SEC13/31A and SEC23A/24D were purified using immobilized metal affinity chromatography from lysates of baculovirus-infected insect cells, as described previously (Kim et al., 2005).

### Cytosol preparation

Plates (20 x 15 cm) of HT-1080 or HT-PC1.1 cells were cultured to confluence, washed with PBS and gently scraped in B88 with protease inhibitors (Roche). Plates (5 x 15 cm) of HEK293T cells were transiently transfected with either 18 μg of empty pEGFP_n1 plasmid, wild-type or mutant KLHL12-3xFLAG in a pcDNA5/FRT/TO vector per 15 cm plate, and cells were collected 24 h after transfection by pipetting with PBS after gently washing with PBS. The collected cells were permeabilized with gentle rocking for 30 min in 80 μg /ml digitonin in 5 ml B88 per 5x 15cm plates with protease inhibitors at concentration suggested by manufacturer (Roche) at 4°C. The cell lysate was collected as a supernatant fraction after centrifugation at 300 xg for 5 min. Digitonin was removed from the cell lysate by incubating with washed Bio-Beads SM2 (dry weight of 1 g beads per 5 x 15 cm dishes) (Bio-Rad) at 4°C overnight. Bio-Beads were removed by centrifugation at 300 xg for 5 min. The supernatant fraction was further centrifuged at 160,000 xg for 30min, and the clarified supernatant was collected and concentrated with an Amicon Ultra-3k (EMD Millipore) filter to 40-80 mg/ml. Small aliquots were frozen in liquid nitrogen and stored at -80°C for future use in vesicle budding reactions.

### Collagenase protection assay

Top fractions collected after Optiprep gradient flotation were pooled and redistributed to each reaction to ensure all samples had the equal starting material. Samples were mixed with or without 0.1 U/μl collagenase (Sigma-Aldrich) in the presence or absence of 1% Triton X-100 (Sigma-Aldrich) and incubated at 0°C or 30°C for 10 min.

### Structured Illumination Microscopy Imaging

Donor ER membrane isolated from cells that expressed PC-GFP was incubated with purified Alexa Fluor 647-conjugated SEC23A/24D and other components, as specified, and budded vesicles were isolated from the top of the flotation gradient as described in "Vesicle budding reaction". Budded vesicles were mounted in Prolong Diamond (Invitrogen) under No.1.5H coverslips (Carl Zeiss AG, Oberkochen, Germany) and set for at least 5 d before imaging to eliminate drift. Z-stacks of 1-2 um were collected in 3 rotations and 5 phases using the 100x/1.46 objective on Elyra PS.1 super-resolution microscope (Carl Zeiss AG) with less than 2% laser power and less than 400 ms exposure in all channels, which was determined using no fluorescence and single channel fluorescence controls. Images were reconstructed and channels were aligned using ZEN Black software (Carl Zeiss AG). 3D iso-surface renderings of each channel were done using Imaris v8.1 (Bitplane USA). Image acquisition, processing, 3D rendering were done using equipment and software in the CNR Biological Imaging Facility at UC Berkeley.

### Flow cytometry

Vesicle budding reactions were scaled up 6x for flow cytometry analyses. After a 7000 xg, 10 min centrifugation of a 600 μl cell-free reaction, 540 μl supernatant was collected from the top. The lipid dye FM4-64 (Thermo Fischer Scientific) was added to the supernatant at 5μg/ml immediately before the flow cytometry analyses. Particles (100,000) were collected for each sample. FSC-A and FSC-H were used to gate for single particles (singlets), which were used for further analysis. Gating of each fluorescent channel was determined by comparing a control sample without any fluorescent labeling and a control that was labeled in a single channel. Data was collected on a BD LSR Fortessa (BD) and analyzed by FlowJo software (FlowJo, LLC Asland, OR, USA). Instrument and software were provided by the LKS flow core facility at UC Berkeley.

### Nanoparticle Tracking Analysis

Size of vesicles budded in vitro were estimated using the NanoSight NS300 instrument equipped with a 405 nm laser (Malvern Instruments Inc., Malvern UK). The Qdot525 positive particles were analyzed with a 500nm filter (fluorescent mode); non-fluorescent particles were analyzed in the scatter mode without a filter. Silica 100 nm microspheres (Polysciences, Inc., Warminster, PA, USA) were analyzed to check instrument performance and used to determine the viscosity coefficient of B88. Vesicles were collected from the top 20 μl of the flotation gradient as described in “vesicle budding reaction” and diluted 40 x with 780 μl filtered B88 (0.02 um, Whatman plc, Maidstone, UK). To label COPII with the fluorescent Qdot525, the 7000 xg supernatant fraction was incubated with mouse anti SEC31A primary antibody (1:100, BD) and Donkey anti mouse IgG secondary antibody Qdot525 conjugate (1:100, Thermo Fisher Scientific) for 1 h at RT. As a labeling control, primary antibody was omitted. The labeled supernatant were used as inputs for flotation (see details in the “vesicle budding reaction”) and 2 x top 20 μl of floated fractions was collected from duplicates, to double the labeling material, and diluted 20 x with 760 μl filtered B88 (0.02 um, Whatman plc). The samples were automatically introduced into the sample chamber at a constant flow rate of 50 (arbitrary manufacturer unit, about 10 μl/min) during 5 repeats of 60 sec captures at camera level 13 in scatter mode and level 15 in fluorescent mode with the Nanosight NTA 3.1 software (Malvern Instruments Inc.). Each sample was measured at 2 different positions of the syringe introduction to minimize the heterogeneous flow of vesicles. The particle size was estimated with detection threshold 3 for fluorescent mode and 5 for scatter mode using the Nanosight NTA 3.1 software, after which “experiment summary” and “particle data” were exported. Particle numbers in each size category was calculated from the “particle data”, where “True” particles with “track length” >3 were pooled, binned, and counted by Excel (Microsoft, Seattle, USA). GraphPad Prism 7 (GraphPad Software, Inc La Jolla, CA, USA) was used for graphing and statistical analyses.

### Electron microscopy of budded vesicles

For morphological analysis of the budded vesicles, the top 80 μl of a supernatant fraction following the 7000 xg spin for 10 min of a 100 μl reaction was fixed with 2% PFA, 0.2% glutaraldehyde in B88 buffer for 15 min at at 4°C and centrifuged at 25,000 xg (Beckman TLS-55 with adaptors for 11x34 mm tubes) for 30 min with slow acceleration and deceleration on to a 0.5 ml agarose bed (2% low melting point, Sigma). This centrifugation speed was optimized to collect large COPII-coated vesicles. After three washes in B88 buffer, the pellet was embedded in 12% gelatin, cut in small blocks and infiltrated with 2.3 M sucrose in 0.1M phosphate buffer at pH 7.4 overnight at 4°C. The blocks were mounted on pins and stored frozen in liquid nitrogen. Ultrathin cryosections were collected on Formvar and carbon coated nickel grids, poststained with 2 *%* uranyl acetate and imaged at 120 kV using a Tecnai 12 Transmission Electron Microscope (FEI, Eindhoven, Netherlands).

## Online supplementary material

**Video 1:** Z sections of PC1 and SEC31A immunolabeled vesicle by 3D STORM. A series of virtual Z-sections in the xy-plane shows encapsulation of PC1 (cyan) by SEC31A (magenta) in three dimensions. Section thickness: 100 nm; step size: 50 nm; scale bar: 200 nm.

**Video 2:** Large COPII vesicles exhibit movement. KI6 cells were transfected with YFP-tagged SEC31A and induced for KLHL12 expression for 7.5 h. The cells were imaged live on a laser scanning confocal microscope. The playback rate is 6 frames per second. Scale bar: 10 μm

**Video 3**: Live cell imaging of SEC31A-YFP vesicles in the presence of 5μM nocodazole KI6 cells were transfected with YFP-tagged SEC31A and induced for KLHL12 expression for 7.5 h and treated with nocodazole for 30 min. The cells were imaged live on a laser scanning confocal microscope. The playback rate is 6 frames per second. Scale bar: 5 μm

**Video 4**: Large COPII vesicles are separated from the ER. KI6 cells were transfected with YFP-tagged SEC31A (red) and CFP tagged ER marker (green) and induced for KLHL12 expression for 7.5 h. The cells were imaged live on a laser scanning confocal microscope. The playback rate is 10 frames per second. Scale bar: 5 μm

**Video 5**: Live cell imaging of SEC31A-YFP and PC1-CFP vesicles. KI6 cells were transfected with YFP-tagged SEC31A (magenta) and CFP tagged PC1 (green) and induced for KLHL12 expression for 7.5 h. The cells were imaged live on a laser scanning confocal microscope. The playback rate is 6 frames per second. Scale bar: 5 μm

## Abbreviations

BFA: brefeldin A
CLEM: Correlative light electron microscopy
COPII: Coat protein complex II
ERES: ER exit sites
ERGIC: ER Golgi intermediate compartment
PC: procollagen
PC1: procollagen I
PFA: Paraformaldehyde
STORM: Stochastic optical reconstruction microscopy
RT: Room temperature

## Acknowledgements

We thank the staff at UC Berkeley shared facilities including Ann Fisher and Alison Kililea (Tissue culture facility), Steven Ruzin and Denise Schichnes (CNR Biolgoical Imaging facility), Kartoosh Heydari (Flow cytometry facility) and Reena Zalpuri (Electron Microscopy lab). We are grateful for advice and guidance by Manfred Auer (LBNL) and Kent McDonald (Director, Electron Microscopy Lab). Research reported in this publication was performed in part at CRL Molecular Imaging Center and The CNR Biological Imaging facility, supported in part by the Gordon and Betty Moore Foundation and The National Institutes of Health S10 program under award numbers 1S10RR026866-01 and 1S10OD018136-01. We also thank past and present members of the Schekman Lab in particular Kanika Bajaj, Yusong Guo, Liang Ge and David Melville. RS is supported as an Investigator of the Howard Hughes Medical Institute and the UC Berkeley Miller Institute of Science. LY was supported in part by the Tang family fellowship. AG was supported in part by the Employee Development Program at Berkeley Lab. S.K. and K.X. acknowledge support from NSF under CHE-1554717, the Pew Biomedical Scholars Award, and the Sloan Research Fellowship. The authors declare no competing financial interests.

## Author contributions

Experiments were designed and interpreted by L.Y., A.G., K.X. and R.S.; A.G. and L.Y. identified the monoclonal antibodies for collagen 1 and performed the immunofluorescence assays; A.G. performed EM and live cell imaging; S.K., performed and analyzed STORM microscopy data; S.B. was instrumental in the initial establishment of the in-vitro reconstitution assay; L.Y. performed and analyzed data from in vitro budding experiments. A.G., L.Y., and R.S. prepared the manuscript.

## Supplementary figures

**Figure S1.**
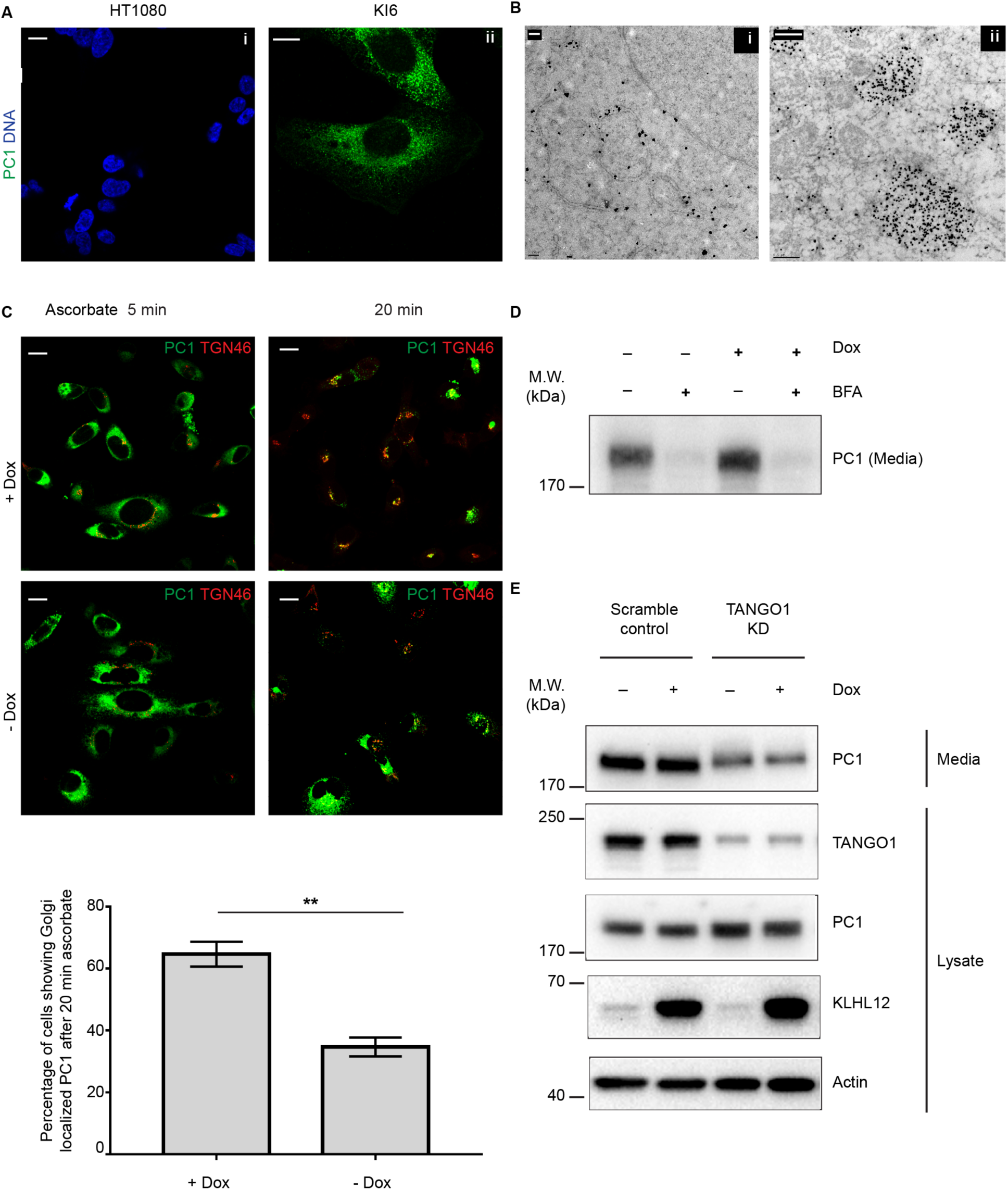
Generation of human fibrosarcoma (KI6) cell line stably transfected with PC1 and doxycycline inducible KLHL12. (A) No PC1 immunofluorescence signal was observed in parent cell line (HT1080) (i), while a uniform reticular pattern was observed in KI6 cells (ii). (B) Pre-embedding immunogold labeling using PC1 antibody in KI6 cells showing reticular pattern in (i) and in large vesicles (ii). (C) KI6 cells treated with doxycycline for 7.5 h (+Dox) and compared with cells without doxycycline (-Dox) after double-immunostaining with PC1 and Golgi marker TGN46 and ascorbate treatments of 5 min and 20 min, respectively. Quantification below shows fraction of cells of a random sampling of 150 cells showing Golgi-localized PC1 in induced versus uninduced cells. Error bar: SD, n=3. (D) Immunoblotting analysis of PC1 in the pre-cleared culture medium secreted by KI6 cells with and without doxycycline induction in the presence or absence of BFA for 8 h. (E) Scramble or TANGO1 siRNA was transfected to KI6 cells for 72 h, and KLHL12 overexpression was induced during the last 7.5 h of knockdown. Pre-cleared culture medium and lysates were collected for immunoblotting analyses. PC1 secretion was inhibited after TANGO1 knockdown in KI6 cells. Scale bars in (A and C): 5 μm, (Bi): 0.2 μm, (Bii): 0.5 μm

**Figure S2:**
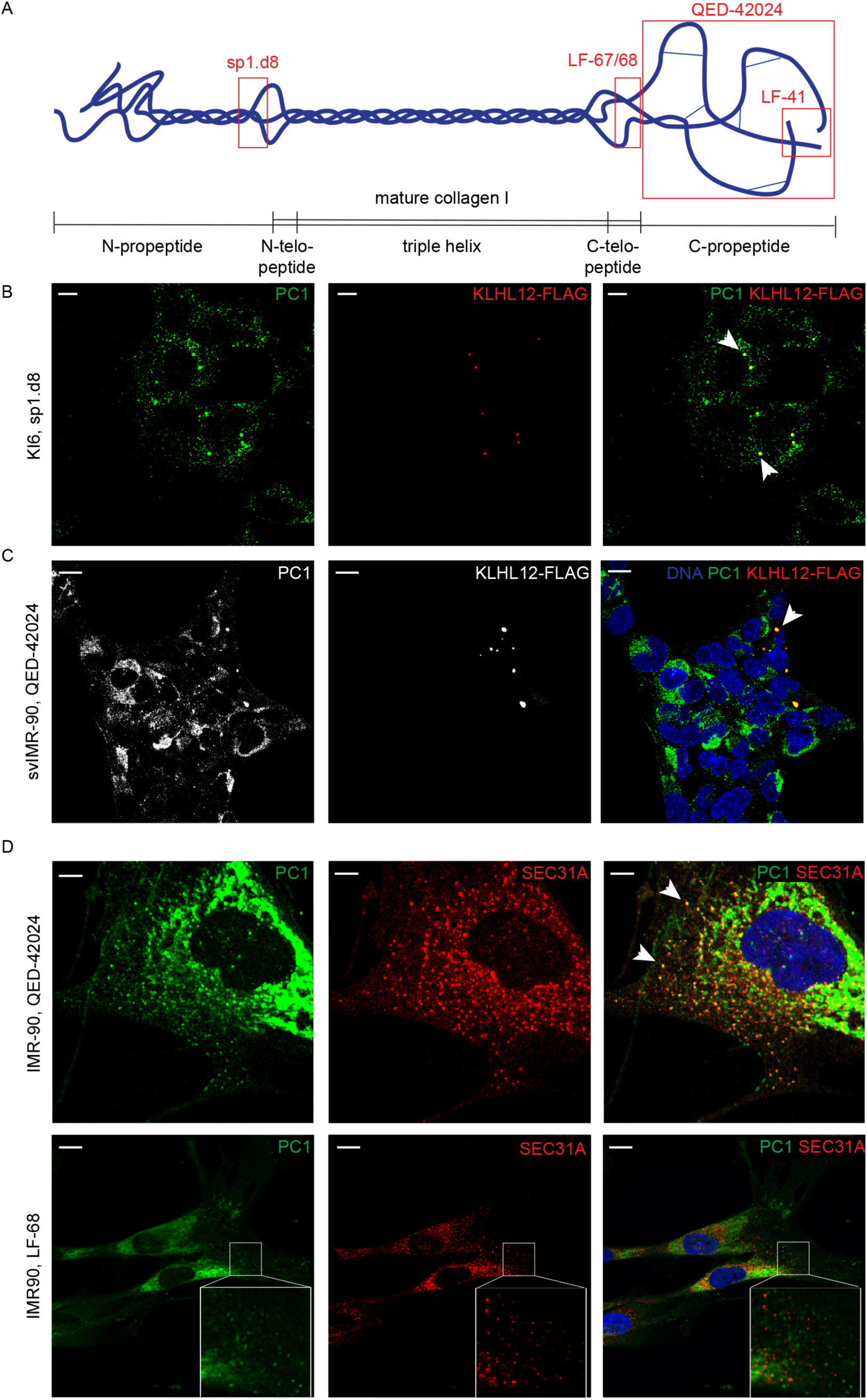
Comparative IF analysis using different PC1 antibodies. (A) Schematic illustration shows PC1 domain structure and sites recognized by different PC1 antibodies discussed in this report (not drawn to scale). Mature collagen I is generated after N and C-propeptides are cleaved from PC1, and the non-triple-helical ends of the mature collagen I are termed N and C-telopeptides. Inter-chain disulfide bonds between C-propeptides are important for trimerization. LF-67 and LF-68 were immunized with the same synthetic peptide sequence of the C-telopeptide and they were used in Jin et al., 2012 and Mironov et al., 2003 respectively. QED-42024 and sp1.d8 are monoclonal antibodies targeting C or N-propeptides respectively. LF-41 was made from synthetic peptide of the end of C-propeptides, and it was exclusively used to label PC1 in all immunoblotting and none of the immunofluorescent experiments in this study. (B) KI6 cells were induced for KLHL12 expression for 7.5 h with doxycycline and labeled with the monoclonal PC1 antibody sp1.d8. Arrows indicate puncta co-localizing for Klhl12 and PC1. (C) svIMR-90 cells were transfected with KLHL12-FLAG. Double immunofluorescence using PC1(QED-42024) and FLAG antibodies revealed puncta that co-localized for both markers (indicated by arrow). (D) Panel shows comparison of labeling using two PC1 antibodies: QED-42024 and LF-68. Arrow indicates SEC31A/PC1 co-localizing puncta. Scale bars in (B and D): 5 μm; (C and E): 10 μm

**Fig S3.**
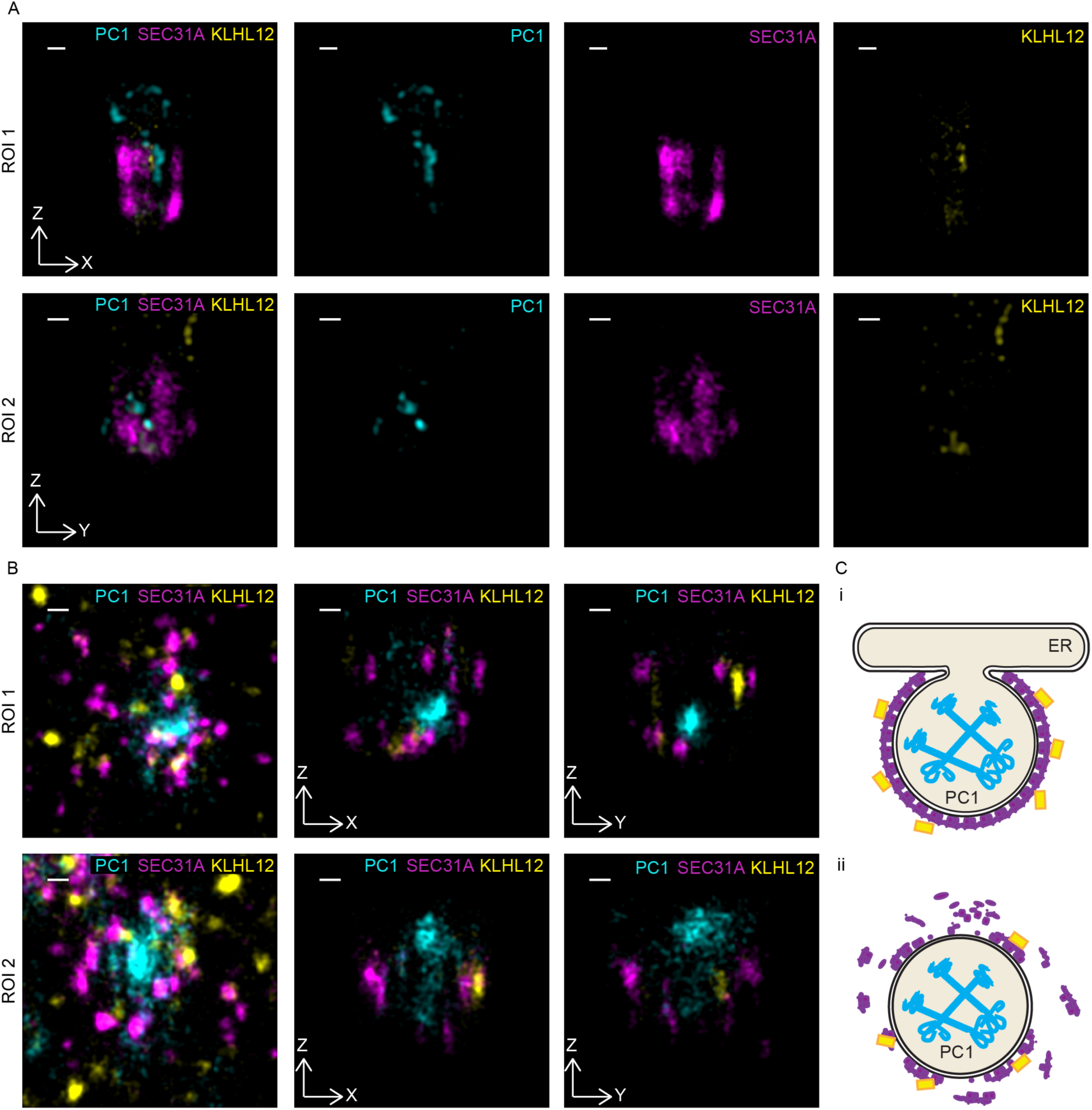
Large partially COPII-coated PC1 carriers observed by STORM. Saos-2 cells were grown at steady state and immunostained for PC1 (cyan), SEC31A (magenta) and KLHL12 (yellow). (A) Two examples of half coated spheres are shown in magnified virtual crosssections. (B) Two examples of PC1 surrounded by patchy coats in XY maximum projection, and virtual cross-sections in XZ and YZ planes in Saos-2 cells. (C) Schematic illustration of nascent budding events (i) observed in (A), and COPII vesicles in the process of decoating (ii) observed in (B). PC1 is drawn in blue, COPII coats in magenta, and KLHL12 in yellow. Scale bars: 100 nm

**Fig S4.**
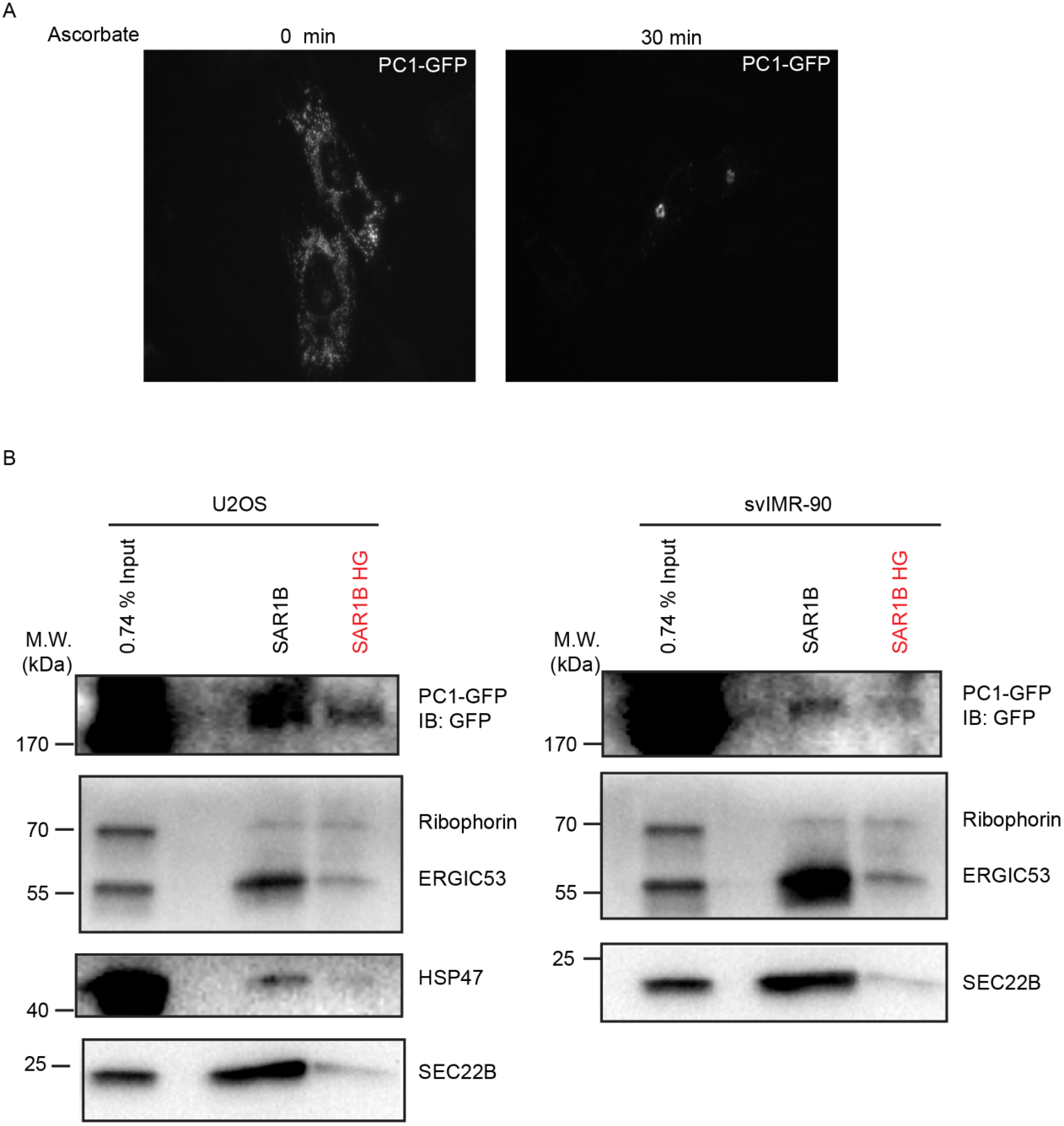
PC1-GFP exits the ER in cells and in the vesicle budding assay. (A) Localization of PC1-GFP was examined in Saos-2 cells transiently transfected with PC1-GFP. The GFP signal localized to the Golgi apparatus af**t**er 30 min ascorbate treatment. (B) Cell-free reaction using donor membrane harvested from cells that overexpressed PC1-GFP and GFP signal at the expected size was detected to exit the ER in a COPII dependent manner.

**Figure S5.**
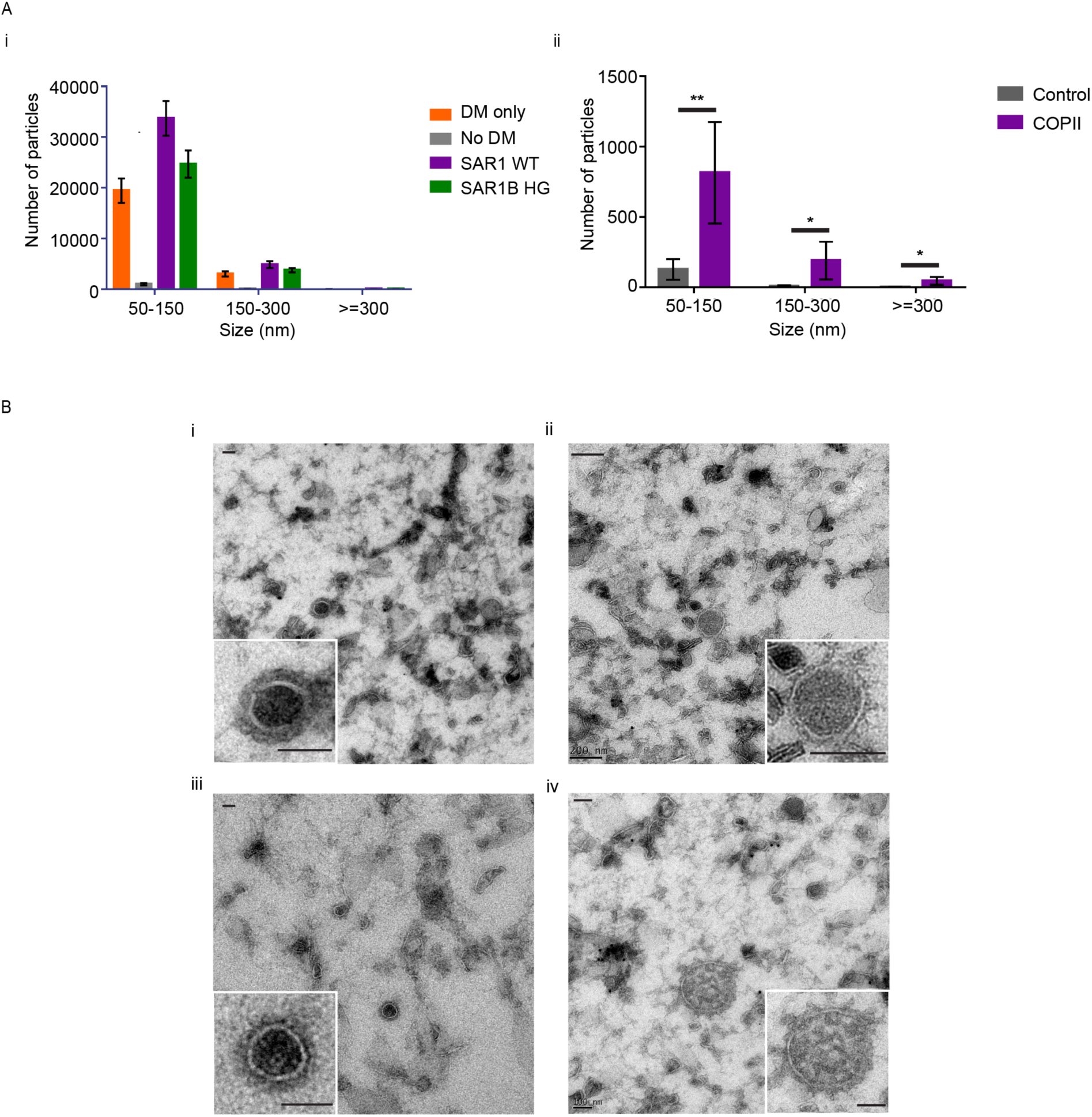
Size distribution of vesicles in the floated fraction. (Ai) Vesicle sizes were measured by nanoparticle tracking analysis (NTA). Bar graphs represent the average number of particles in each size category (n=3, Error bar: SD). Budding reactions with donor membrane, nucleotides, recombinant COPII, and cytosol (“Everything”, purple) was used as positive control. Reactions using only donor membrane (“DM only”, orange), everything but the donor membrane (“No DM”, gray), and everything but SAR1B H79G instead of wild-type SAR1B (“HG control”, green) were negative controls. The donor membrane itself (orange) contributed an averaged 57.2% (SD=2.64%, n=3) of all particles detected in the positive control (purple). An average of 27.0% (SD=1.36%, n=3) particles detected in the budding reaction were COPII vesicles, deduced from the differences between the positive and HG controls. (Aii) Size distribution of COPII carriers in the floated fraction quantified by fluorescent NTA to overcome high background of non-COPII particles. COPII carriers were labeled with mouse anti-SEC31A primary antibody and Qdot525 conjugated anti-mouse IgG secondary antibodies (Purple). Controls were incubated without primary antibodies (Gray). Bar graphs represent the average number of fluorescent particles in each size category and the antibody labeling of COPII vesicles was successful shown by the statistical significance compared with the controls (n=5, Error bar: SD, paired t-test, p=0.0094, 0.0346, 0.0242 for categories 50-150 nm, 150-300 nm, and >=300 nm respectively). COPII vesicles in the range of 50-150 nm were about 18 times more enriched on average than large COPII that are 300-1000 nm. The amount of conventional COPII estimated by fluorescence NTA appears to under-represent the amount deduced from scatter NTA in (Ai), possibly due to the dynamic shedding of the COPII coat and limitation of antibody affinities. The intensity of luminal GFP was too weak to be detected by fluorescent NTA. Due to this technical limitation, we were not able to quantify the size distribution of PC1-carrying COPII by NTA. (B) Morphology of membrane vesicles generated from *in vitro* budding reaction (i-iv) Ultrathin cryosections of sample collected from an *in vitro* budding reaction show an overview of differently sized membrane vesicles and an inset showing a magnified view of a single vesicle (i) coated ~150 nm membrane vesicle (ii) uncoated or partially coated ~200 nm vesicle (iii) coated ~225 nm membrane vesicle and (iv) ~300 nm vesicle. Scale bars in (i and iv, including insets): 100 nm; (ii and iii, including insets): 200 nm.

